# Bridging brain and cognition: A multilayer network analysis of brain structural covariance and general intelligence in a developmental sample of struggling learners

**DOI:** 10.1101/2020.11.15.383869

**Authors:** Ivan L. Simpson-Kent, Eiko I. Fried, Danyal Akarca, Silvana Mareva, Edward T. Bullmore, the CALM Team, Rogier A. Kievit

## Abstract

Network analytic methods that are ubiquitous in other areas, such as systems neuroscience, have recently been used to test network theories in psychology, including intelligence research. The network or mutualism theory of intelligence proposes that the statistical associations among cognitive abilities (e.g., specific abilities such as vocabulary or memory) stem from causal relations among them throughout development. In this study, we used network models (specifically LASSO) of cognitive abilities and brain structural covariance (grey and white matter) to simultaneously model brain-behavior relationships essential for general intelligence in a large (behavioral, N=805; cortical volume, N=246; fractional anisotropy, N=165), developmental (ages 5-18) cohort of struggling learners (CALM). We found that mostly positive, small partial correlations pervade our cognitive, neural, and multilayer networks. Moreover, using community detection (Walktrap algorithm) and calculating node centrality (absolute strength and bridge strength), we found convergent evidence that subsets of both cognitive and neural nodes play an intermediary role ‘between’ brain and behavior. We discuss implications and possible avenues for future studies.

## INTRODUCTION

General intelligence or *g* (Spearman 1904) captures cognitive ability across a variety of domains and predicts a wide range of important life outcomes such as educational and occupational achievement (Hegelund et al. 2018), and mortality (Calvin et al. 2011). As individual differences in intelligence found in childhood remain surprisingly stable across the lifespan (Deary et al. 2004), a deeper understanding of the mechanisms of cognitive ability is crucial to supporting individuals, especially those considered ‘low-performing’ (e.g., students struggling to learn in school). Recent work in network analysis has shed new light on both the cognitive abilities that make up general intelligence (Kievit, Hofman, and Nation 2019; van der Maas et al. 2017), as well as the brain systems purported to support them (Girn, Mills, and Christoff 2019; Seidlitz et al. 2018).

In psychological sciences, factor models have traditionally been used to study intelligence (e.g., Carroll 1993). However, in the last two decades there has been a rise in use of the statistical tools of network science (Barabási 2016) to show that examining relationships between cognitive abilities can help us better understand the *development* of general intelligence. For instance, the mutualism model (van der Maas et al. 2006) was inspired by an ecosystem model of prey-predator relations, and states that the positive manifold (Spearman 1904) *emerges* gradually from the positive interactions among different cognitive abilities (i.e., reasoning and vocabulary) over time (see Kievit et al. 2017; Kievit, Hofman, and Nation 2019). Hence, *g* can arise even from originally weakly correlated cognitive faculties. This highlights the need to both conceptualize traits, abilities, or psychological constructs such as general intelligence as complex dynamical systems, as well as use appropriate statistical models (i.e., network analysis) to estimate relationships among elements of the systems under investigation (Fried 2020a; Fried and Robinaugh 2020).

Although longitudinal data are preferable to study developmental questions (e.g., Kievit et al. 2013; Rhemtulla, Bork, and Cramer 2020), cross-sectional network analysis can provide important insights. Recent innovations in network psychometrics (Epskamp et al. 2018) have led to a rapid increase in popularity of behavioral network analysis, especially in psychopathology (Borsboom 2017; Robinaugh et al. 2019). In this framework, psychological constructs are theorized as complex systems and relationships (edges) between nodes (e.g., item responses on a questionnaire) are estimated using weighted partial correlation networks, which enable determination of conditional dependencies among variables after controlling for the associations among every other node in the network (Epskamp et al. 2018).

Recently, this approach has also been used to analyze cross-sectional data on general intelligence. For instance, both Kan, van der Maas, and Levine 2019 (N=1,800; age range: 16-89 years) as well as Schmank et al. 2019 (N=1,112; age range: 12-90 years) found support for mutualism in the WAIS-IV cognitive battery (Wechsler 2008), such that a network model showed better fit to the pattern of intelligence scores compared to a latent variable approach (*g* factor). Lastly, Mareva and Holmes 2020, in two separate samples, one the same group of struggling learners as studied here (see Holmes et al. 2019 for detailed overview of the cohort) but with less participants (N=350), no neuroimaging data, and including tasks not analyzed in this study (e.g., motor speed and tower achievement), observed links between cognitive abilities and learning, especially between mathematics skills and more “domain-general” faculties such as backward digit span and matrix reasoning.

In neuroscience, network analysis methods have been widely used to describe the relations among brain regions, ushering in the field of network neuroscience (Bassett and Sporns 2017; Fornito, Zalesky, and Bullmore 2016). Rather than focusing on individual brain regions in isolation, the brain is conceived as a complex system of interconnected networks that facilitate behavioral functions ranging from sensorimotor control to learning. In this light, several influential studies have revealed pervasive properties of brain networks such as small-world topology (Bassett and Bullmore 2006; Bassett and Bullmore 2017), modularity (Meunier, Lambiotte, and Bullmore 2010; Sporns and Betzel 2016) and ‘rich-club’ connector hubs (Heuvel and Sporns 2011), consistent with an economical trade-off between minimizing wiring cost and maximizing efficiency (e.g., information transfer) that enable adaptive behavior (Bullmore and Sporns 2012).

One proposal attempting to explain general intelligence using network neuroscience is The Network Neuroscience Theory of Human Intelligence (NNTHI, Barbey 2018). Barbey argues that general intelligence arises from the dynamic small-world typology of the brain, which permits transitions between “regular” or “easy-to-reach” network states (needed to access prior knowledge for specific abilities) and “random” or “difficult-to-reach” (required to integrate information for broad abilities) network states (i.e., as in network control theory, see Gu et al. 2015). Together, this constrained flexibility allows the brain to adapt to novel cognitive domains (e.g., in abstract reasoning) while still preserving access to previously learned skills (e.g., from schooling).

Evidence supporting the NNTHI has been inconclusive so far (Girn, Mills, and Christoff 2019). However, two recent studies, although not directly testing the NNTHI, have shed light on the network neuroscience of cognition. Bertolero et al. 2018 found that a mechanistic model assuming that “connector hubs” (diverse club nodes, see Bertolero, Yeo, and D’Esposito 2017), which regulate the activity of their neighboring communities to be more modular but maintain the capability of “task appropriate information integration across communities”, significantly predicted higher cognitive performance on various tasks including language and working memory. Furthermore, in the same sample studied here, Akarca et al. 2020 applied a generative network modelling approach to simulate the growth of brain network connectomes, finding that it is possible to simulate structural networks with statistical properties mirroring the spatially embedding of those observed. The parameters of these generative models were shown to correlate with neuroimaging measures not used to train the models (including grey matter measures), cognitive performance (including vocabulary and mathematics) and relate to gene expression in the cortex. Together these studies point the field toward a better mechanistic understanding of the development of human brain structure, function, and their relationship with cognitive ability.

Although network approaches have provided unique insights within cognitive neuroscience as well as cognitive psychology, few studies have integrated them into a so-called multilayer network paradigm (Bianconi 2018), which models the relationships among variables simultaneously across time (e.g., days, weeks, months, and years) and/or levels of organization (e.g., behavior and brain variables). Two studies have recently pushed this boundary. Hilland et al. 2020 examined the relations between brain structure (cortical thickness and volume) and depression symptoms. They found (via a partial correlation network model) that certain clusters of brain regions (cingulate, fusiform gyrus, hippocampus, and insula) were conditionally dependent with a subset of depression symptoms (crying, irritability and sadness). Secondly, in 172 male autistic participants (ages 10-21 years), Bathelt, Geurts, and Borsboom 2020 used “network-based regression” to estimate the relationship between the unique variance of both the autism symptom network and functional brain connectivity (resting-state fMRI). Moreover, they applied Bayesian network analysis to create a directed acyclic graph between autism symptoms subscores and their neural correlates. They found that communication and social behavior were predicted by their respective resting-state MRI neural correlates (termed Comm Brain and Social Brain).

This study builds on these findings and the recent studies mentioned above, by combining a network psychometrics approach to understand individual differences in cognitive ability (general intelligence) with structural covariance networks derived from structural brain properties (grey matter cortical volume and white matter fractional anisotropy). Doing so, we created a *network of networks*, which differs from multiplex (same nodes, different edge types across layers) and multi-slice (same nodes and edge types over time such as in fMRI time-series data) networks (see figure 4.1 of Bianconi 2018). The advantages of applying this approach are threefold and complementary. First, it places the brain and behavior, which often do not map onto each other in a simple and reductionistic one-to-one fashion, into the same analytical paradigm (network analysis). This allows for simultaneous estimations and easier visualizations of potential causal links between cognition and structural brain properties, which to our knowledge, has only been done in a similar way in two other studies, one involving depression (Hilland et al. 2020), the other in autism (Bathelt, Geurts, and Borsboom 2020). Second, it enables the use of community detection algorithms to tease apart major clusters of cognitive abilities, which could help pinpoint potential intervention targets (e.g., using cognitive training and/or transcranial magnetic stimulation). Lastly, it aids in establishing a coherent framework for theory building, which has been lacking in both the neuroscience (Levenstein et al. 2020) and psychological (Fried 2020a) literature, by treating both the brain (algorithmic) and behavior (computational) as *equally important* levels of analysis to study (Marr and Poggio 1976), and attempting to more directly translate findings from one level to the other. Ultimately, the hope is that relations between brain-behavior nodes can help identify candidate targets (e.g., nodes that bridge the brain and cognition) for future interventions in developmental samples of struggling learners.

In this study, we used network models (specifically LASSO) of cognitive abilities and brain structural covariance (grey and white matter) to simultaneously model brain-behavior relationships essential for general intelligence in a large (behavioral, N=805; cortical volume, N=246; fractional anisotropy, N=165), developmental (ages 5-18) cohort of struggling learners (CALM). We found that mostly positive, small partial correlations pervade our cognitive, neural, and multilayer networks. Moreover, using community detection (Walktrap algorithm) and calculating node centrality (absolute strength and bridge strength), we found convergent evidence that subsets of both cognitive and neural nodes play an intermediary role ‘between’ brain and behavior. We discuss implications and possible avenues for future studies.

## METHODS

### Participants

The present cross-sectional sample (behavioral, N=805; cortical volume, N=246; fractional anisotropy, N=165; age range: 5 to 18 years) was obtained from the Centre for Attention, Learning and Memory (CALM) located in Cambridge, UK (Holmes et al. 2019). This developmental cohort consists of children and adolescents recruited by referrals for perceived difficulties in attention, memory, language, reading and/or mathematics problems. A formal diagnosis was neither required nor an exclusion criterion. Exclusion criteria included any known significant and uncorrected problems in vision or hearing, and/or being a non-native English speaker.

Cognitive data were obtained on a one-to-one basis by an examiner in a designated child-friendly testing room. The tasks analyzed in this study comprised a comprehensive array of standardized assessments of cognitive ability including crystallized intelligence (peabody picture vocabulary test, spelling, single word reading, and numerical operations), fluid intelligence (matrix reasoning), and working memory (forward and backward digit recall, Mr. X, dot matrix, and following instructions). See Table 1 for tasks descriptions, relevant citations, and summary statistics.

**Table 1.**
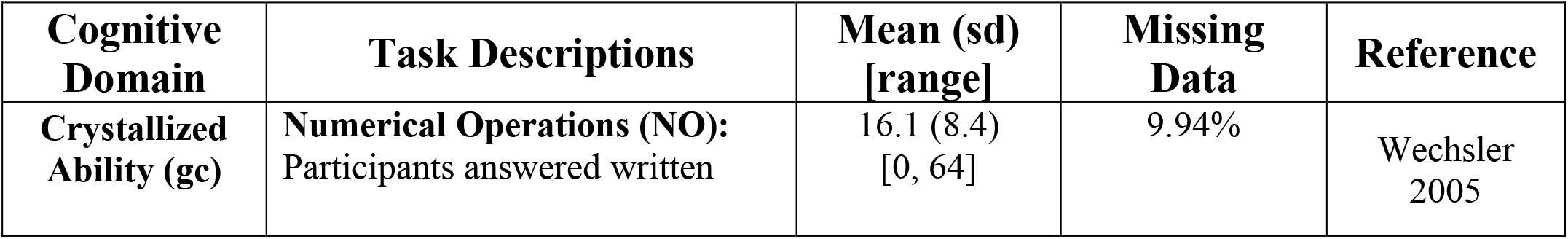
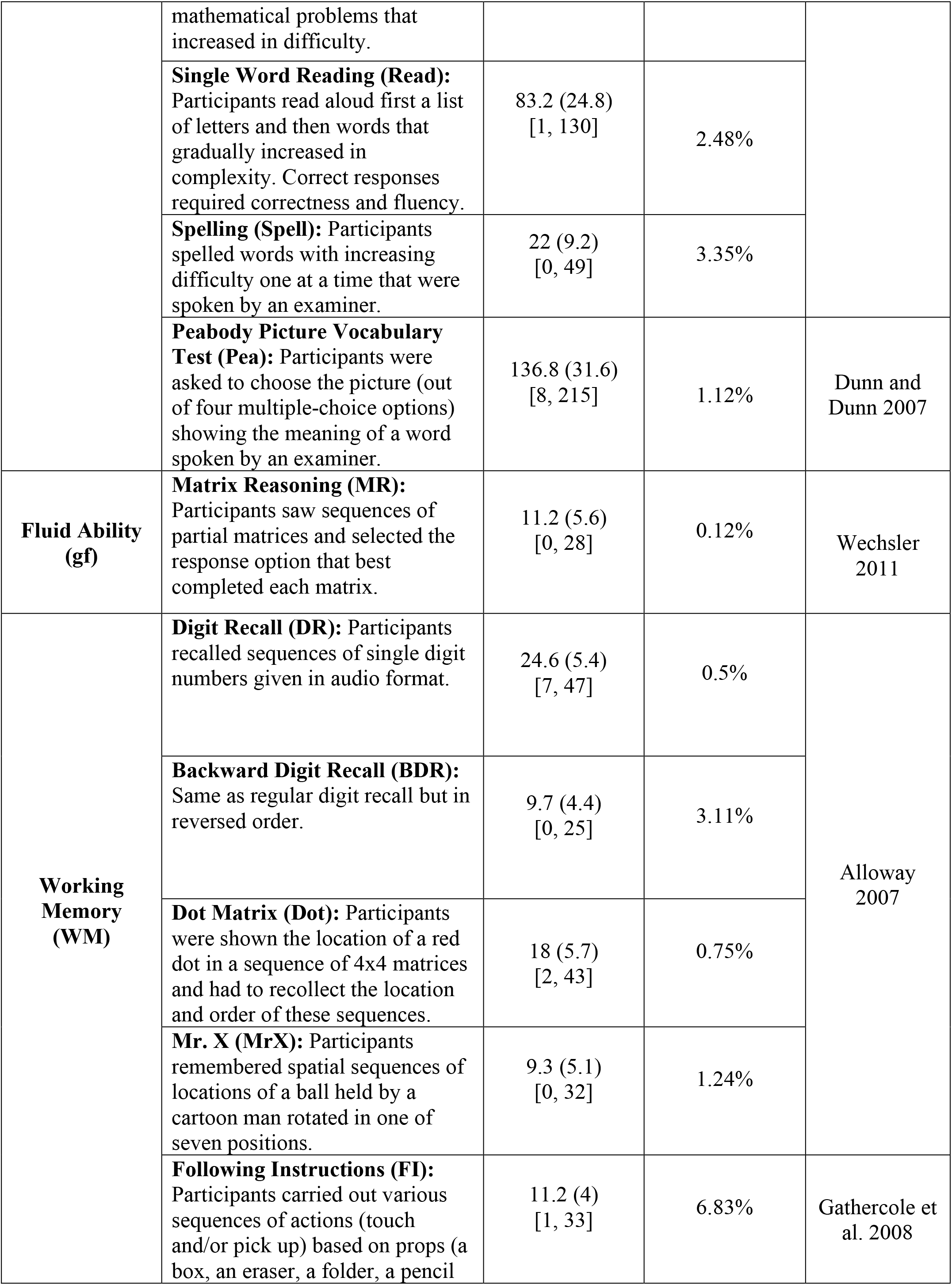
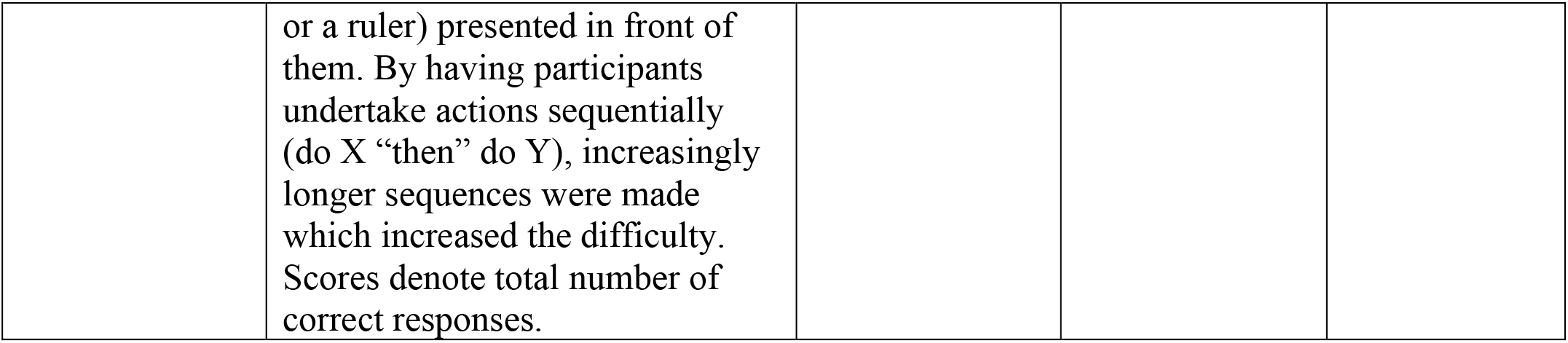
List, descriptions, and summary statistics (mean, standard deviation, range, and percentage of missing data) of cognitive assessments used in the study from the CALM sample. Note, tasks descriptions (except following instructions) are taken directly or paraphrased from Simpson-Kent et al. 2020.

Participants were allotted regular breaks throughout each session. When necessary, testing was split into two separate sessions for participants who did not complete the assessments in a single sitting. A subset of participants also underwent MRI scanning (see below for details).

### Structural Neuroimaging: Cortical Volume (CV) and Fractional Anisotropy (FA)

CALM neuroimaging data were obtained at the MRC Cognition and Brain Sciences Unit, Cambridge, UK. Scans were acquired on the Siemens 3 T Tim Trio system (Siemens Healthcare, Erlangen, Germany) via 32-channel quadrature head coil. T1-weighted volume scans were acquired using a whole brain coverage 3D magnetization-prepared rapid acquisition gradient echo (MPRAGE) sequence, with 1 mm isotropic image resolution. The following parameters were used: Repetition time (TR) = 2250 ms; Echo time (TE) = 3.02 ms; Inversion time (TI) = 900 ms; Flip angle = 9 degrees; Voxel dimensions = 1 mm isotropic; GRAPPA acceleration factor = 2. Diffusion-Weighted Images (DWI) were acquired using a Diffusion Tensor Imaging (DTI) sequence with 64 diffusion gradient directions, with a b-value of 1000 s/mm^2^, plus one image acquired with a b-value of 0. Relevant parameters include: TR = 8500 ms, TE = 90 ms, voxel dimensions = 2 mm isotropic.

We undertook several procedures to ensure adequate MRI data quality and minimize potential biases due to subject movement. For all participants in CALM, children were trained to lie still inside a realistic mock scanner prior to their scan. All T1-weighted images and FA maps were examined by an expert to remove low quality scans. Moreover, only data with a maximum between-volume displacement below 3 mm were included in the analyses.

As our grey matter metric, we use region-based cortical volume (CV in mm^3^, N=246, averaged across contralateral homologues), based on the Desikan-Killiany atlas (Desikan et al. 2006) and defined as the distance between the outer edge of cortical grey matter and subcortical white matter (Bruce Fischl and Dale 2000). Tissue classification and anatomical labelling was performed on the basis of the T1-weighted scan using FreeSurfer v5.3.0 software which is documented and freely available for download online (http://surfer.nmr.mgh.harvard.edu/). The technical details of these procedures are described in prior publications (Dale, Fischl, and Sereno 1999; Fischl, Sereno, and Dale 1999; Fischl et al. 2002). FreeSurfer morphology output statistics were computed for each ROI, and also included cortical thickness and surface area (see Supplementary Material for analyses involving these two metrics). Based on a recent meta-analyses on functional and structural correlates of intelligence (Basten, Hilger, and Fiebach 2015), we included a subset of 10 cortical volume regions in this study: caudal anterior cingulate (CAC), caudal middle frontal gyrus (CMF), frontal pole (FP), medial orbitofrontal cortex (MOF), rostral anterior cingulate gyrus (RAC), rostral middle frontal gyrus (RMF), superior frontal gyrus (SFG), superior temporal gyrus (STG), supramarginal gyrus (SMG), and transverse temporal gyrus (TTG).

From a subset our neuroimaging data (see Simpson-Kent et al. 2020), we also calculated fractional anisotropy (FA, N=165), a proxy for white matter integrity (Wandell 2016). We included 10 regions using the Johns Hopkins University DTI-based white matter tractography atlas (see Hua et al. 2008): anterior thalamic radiations (ATR), corticospinal tract (CST), cingulate gyrus (CING), cingulum [hippocampus] (CINGh), forceps major (FMaj), forceps minor (FMin), inferior fronto-occipital fasciculus (IFOF), inferior longitudinal fasciculus (ILF), superior longitudinal fasciculus (SLF), and uncinate fasciculus (UNC).

All steps to compute regional CV estimation and FA maps were implemented using NiPyPe v0.13.0 (see https://nipype.readthedocs.io/en/latest/). To create a brain mask based on the b0-weighted image (FSL BET; Smith 2002) and correct for movement and eddy current-induced distortions (eddy; Graham, Drobnjak, and Zhang 2016), diffusion-weighted images were pre-processed. The diffusion tensor model was then fitted and fractional anisotropy (FA) maps were calculated using *dtifit*. Images with a between-image displacement >3 mm were then excluded from subsequent analysis steps. This was completed using FSL v5.0.9. To extract FA values for major white matter tracts. FA images were registered to the FMRIB58 FA template in MNI space using a sequence of rigid, affine, and symmetric diffeomorphic image registration (SyN). This was implemented in ANTS v1.9 (Avants et al. 2008). For all participants, visual inspection indicated good image registration. Binary masks from a probabilistic white matter atlas (thresholded at >50 % probability) in the same space were applied to extract FA values.

We used these region based measures to study brain structural covariance (Alexander-Bloch, Giedd, and Bullmore 2013), which have been used in cross-sectional and longitudinal designs of cognitive ability in childhood and adolescence (e.g., Solé-Casals et al. 2019; see Kievit and Simpson-Kent 2021 for a recent review of longitudinal studies). Emerging theoretical proposals emphasize the role of networks of brain areas in producing intelligent behavior (e.g., Parieto-Frontal Integration Theory (P-FIT), Jung and Haier 2007 and The Network Neuroscience Theory of Human Intelligence, Barbey 2018) rather than individual regions-of-interest (ROIs) in isolation (e.g., primarily the prefrontal cortex). As stated above, we selected 10 grey matter and 10 white matter ROIs based upon combined evidence from a recent meta-analysis (Basten, Hilger, and Fiebach 2015) on associations between functional and structural ROIs and cognitive ability that further extended the P-FIT theory, but also more recent work done in two large cohorts, one in longitudinal analysis of the UK Biobank sample (grey matter, Kievit et al. 2018) and another in a the same (cross-sectional) developmental cohort, although with a smaller sample size (cognitive data, N=551; neural data, N=165), studied here (white matter, Simpson-Kent et al. 2020).

For a correlation plot of cognitive tasks and neuroimaging measures, see Figure 1. To view age trends of cognitive tasks and structural neuroimaging measures, see Figure 2. Lastly, see Figure 3 for illustrations of ROIs analyzed in this study.

**Figure 1.**
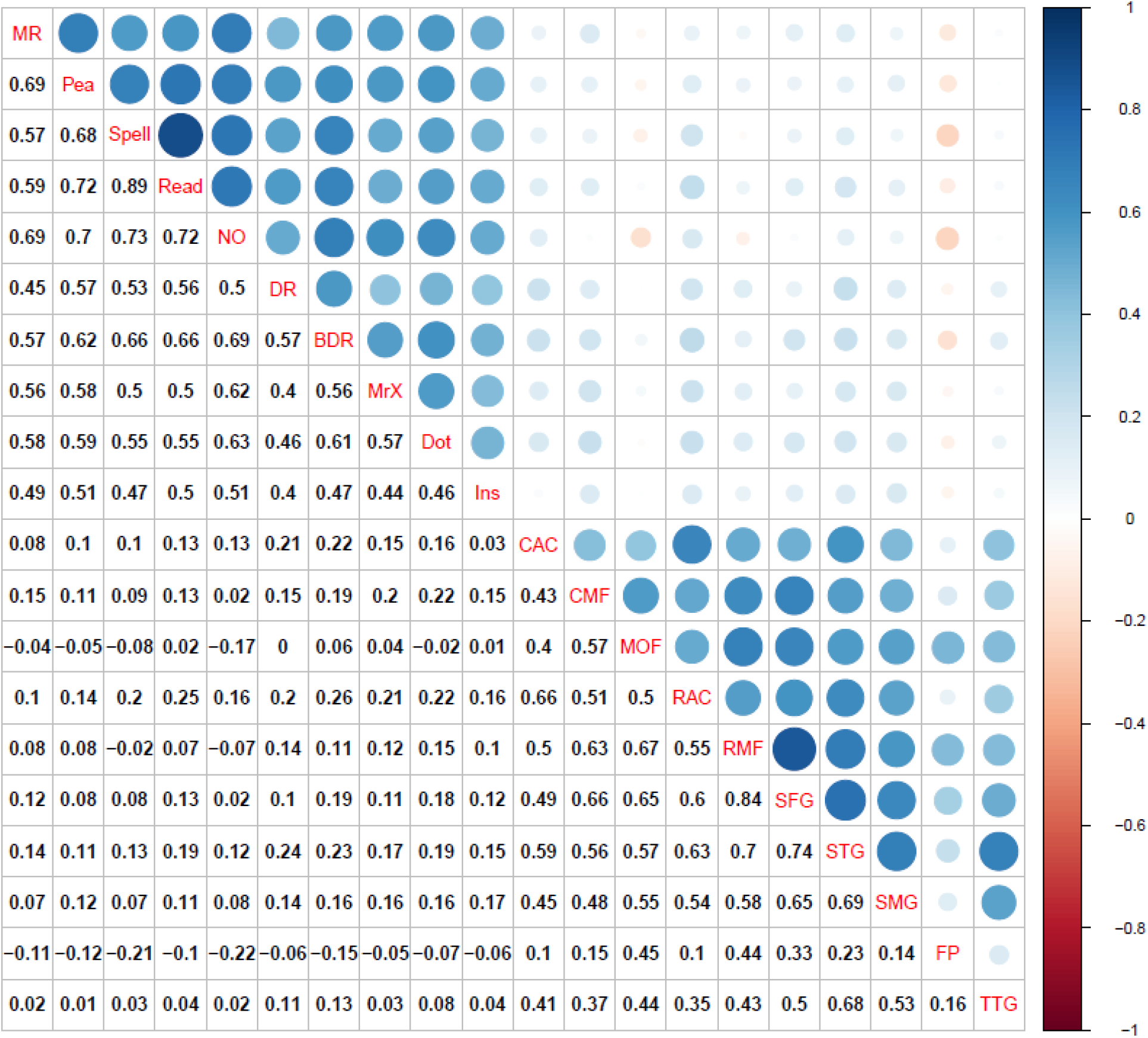
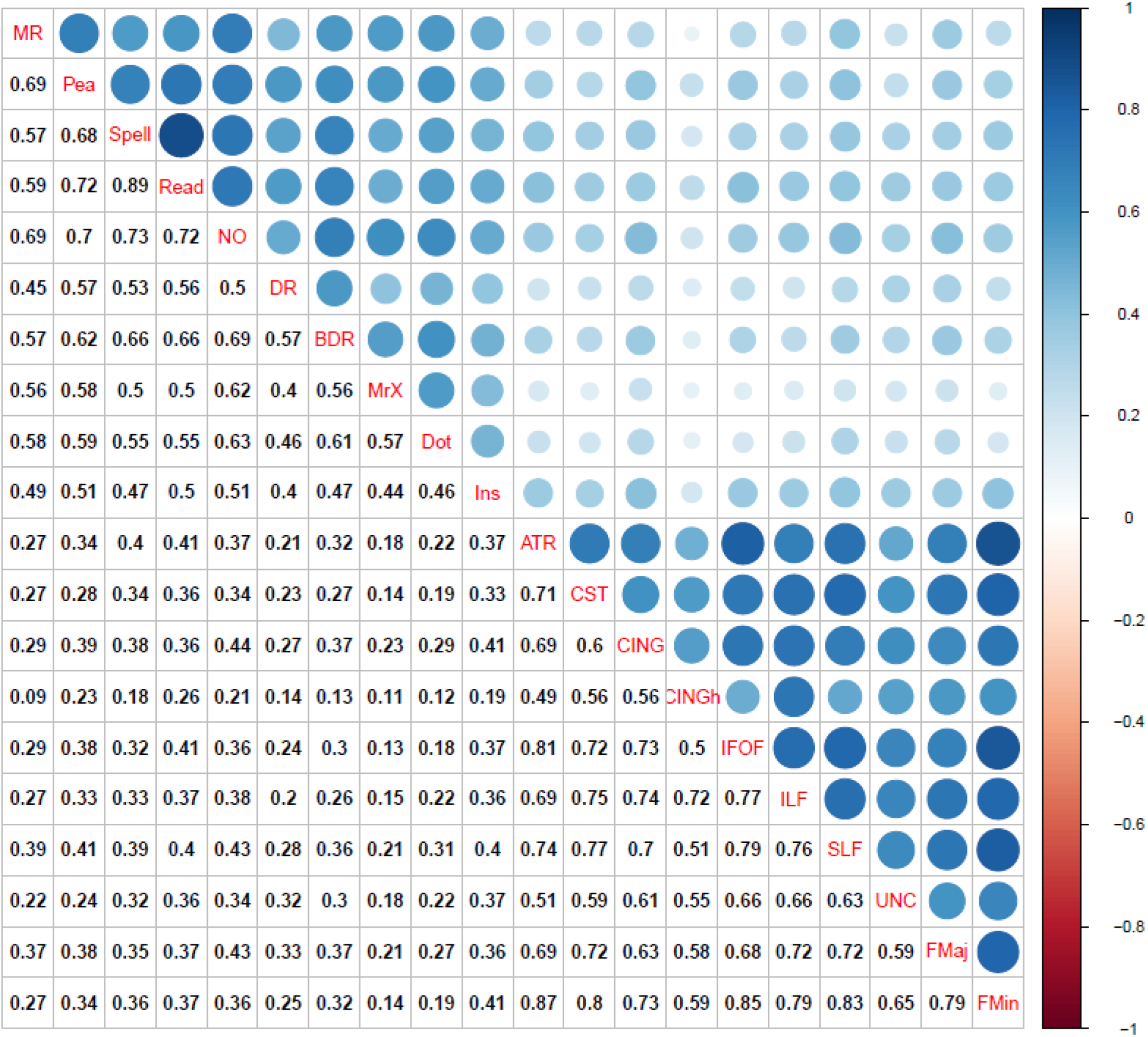
Top: Correlation plot for cognitive raw scores and bilateral cortical volume ROIs. Bottom: Correlation plot for cognitive raw scores and bilateral fractional anisotropy ROIs. All coefficients shown are Pearson correlations. Blue represents positive correlations while red signifies negative correlations among variables. Size of circles indicates the magnitude of the association (e.g., larger circle=higher correlation). Correlations calculated using pairwise complete observations. Abbreviations: matrix reasoning (MR), peabody picture vocabulary test (Pea), Spelling (Spell), single word reading (Read), numerical operations (NO), digit recall (DR), backward digit recall (BDR), Mr. X (MrX), dot matrix (Dot), following instructions (Ins), caudal anterior cingulate (CAC), caudal middle frontal gyrus (CMF), medial orbitofrontal cortex (MOF), rostral anterior cingulate gyrus (RAC), rostral middle frontal gyrus (RMF), superior frontal gyrus (SFG), superior temporal gyrus (STG), supramarginal gyrus (SMG), frontal pole (FP), transverse temporal gyrus (TTG), anterior thalamic radiations (ATR), corticospinal tract (CST), cingulate gyrus (CING), cingulum [hippocampus] (CINGh), inferior fronto-occipital fasciculus (IFOF), inferior longitudinal fasciculus (ILF), superior longitudinal fasciculus (SLF), uncinate fasciculus (UNC), forceps major (FMaj), and forceps minor (FMin).

**Figure 2.**
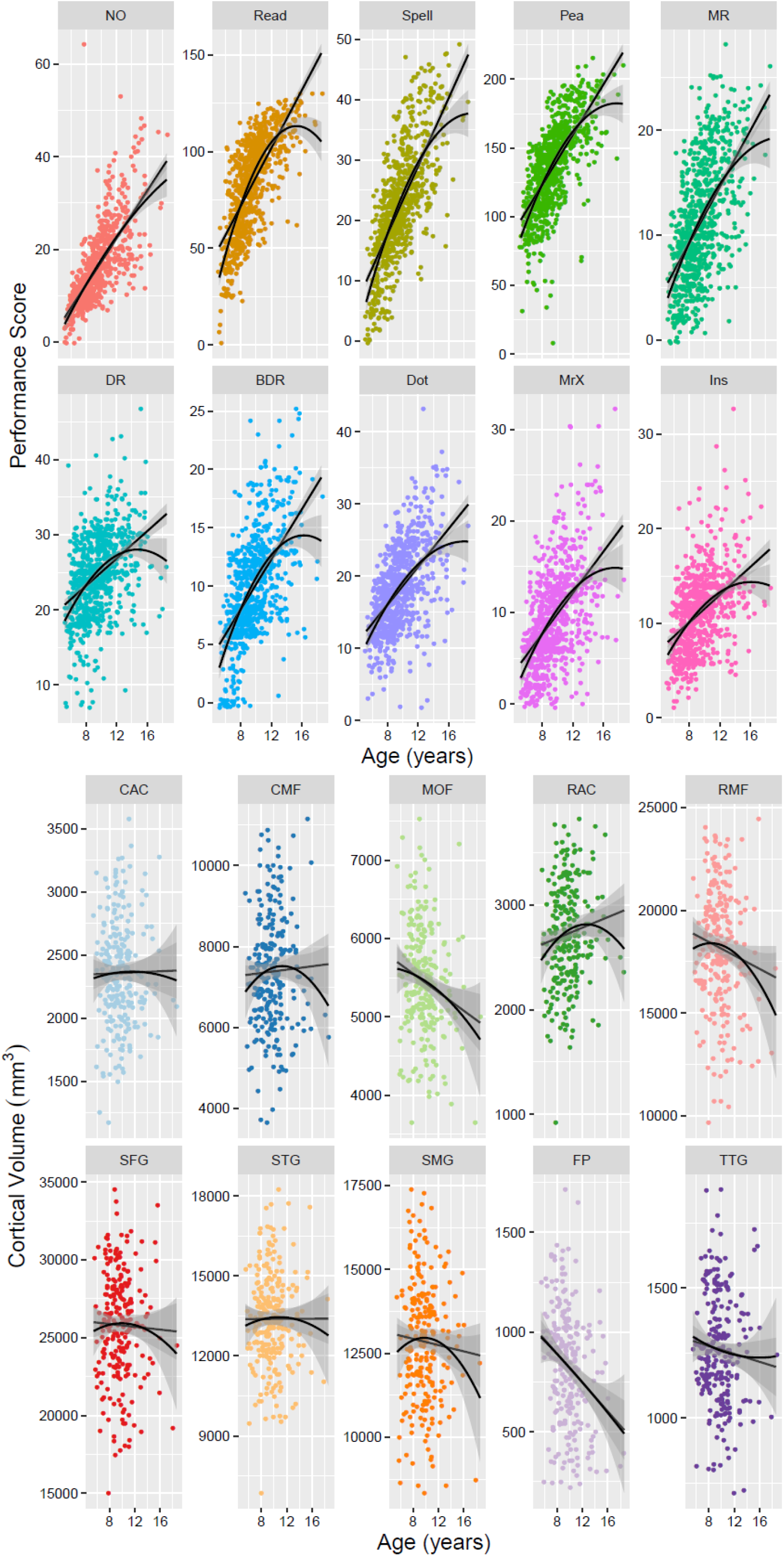
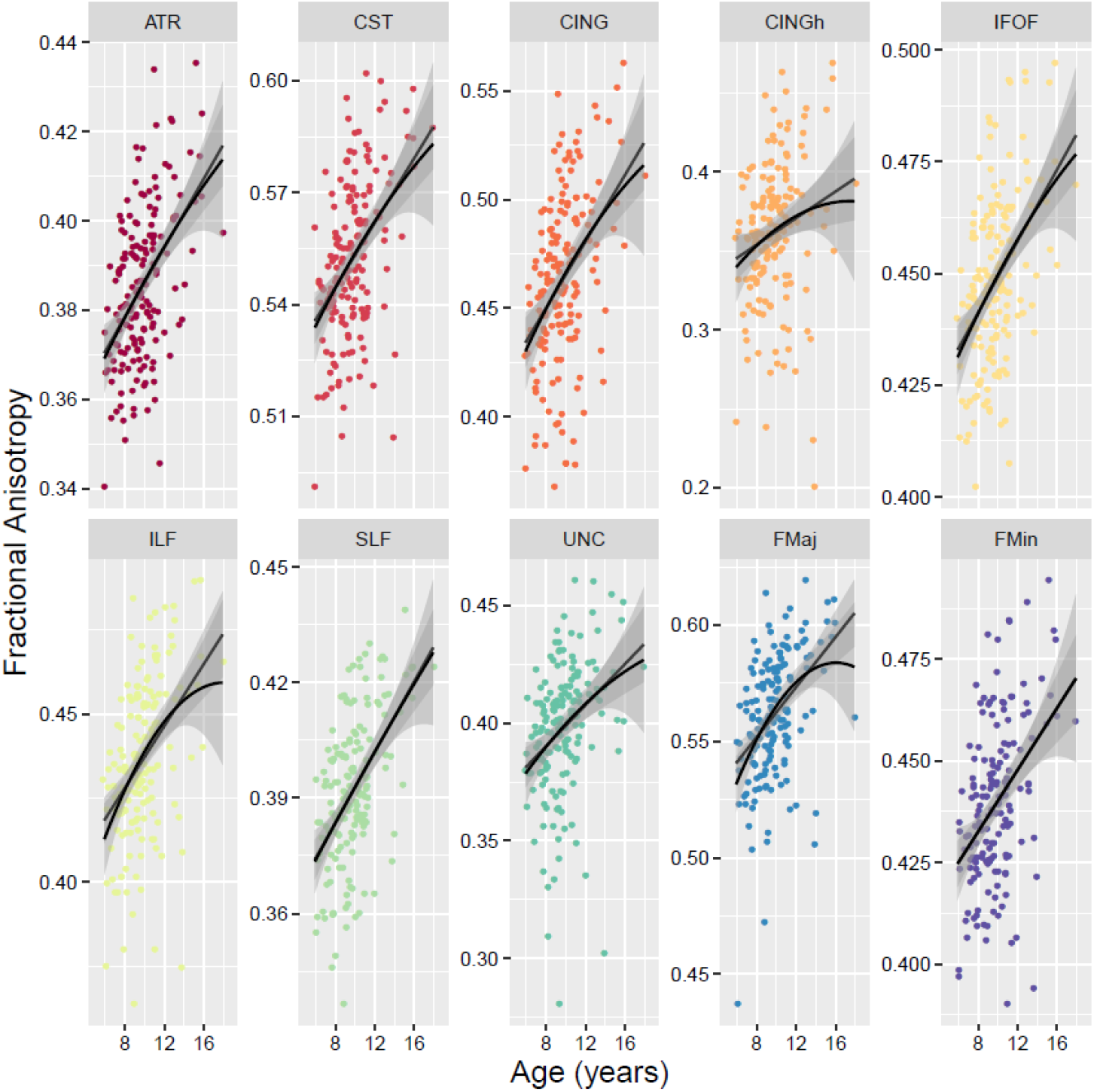
Cross-sectional scatterplots for cognitive raw scores (top), bilateral cortical volume (middle) and bilateral fractional anisotropy (bottom). Solid lines represent linear and polynomial fit while shades indicate 95% confidence intervals. Abbreviations: matrix reasoning (MR), peabody picture vocabulary test (Pea), Spelling (Spell), single word reading (Read), numerical operations (NO), digit recall (DR), backward digit recall (BDR), Mr. X (MrX), dot matrix (Dot), following instructions (Ins), caudal anterior cingulate (CAC), caudal middle frontal gyrus (CMF), medial orbitofrontal cortex (MOF), rostral anterior cingulate gyrus (RAC), rostral middle frontal gyrus (RMF), superior frontal gyrus (SFG), superior temporal gyrus (STG), supramarginal gyrus (SMG), frontal pole (FP), transverse temporal gyrus (TTG), anterior thalamic radiations (ATR), corticospinal tract (CST), cingulate gyrus (CING), cingulum [hippocampus] (CINGh), inferior fronto-occipital fasciculus (IFOF), inferior longitudinal fasciculus (ILF), superior longitudinal fasciculus (SLF), uncinate fasciculus (UNC), forceps major (FMaj), and forceps minor (FMin).

**Figure 3.**
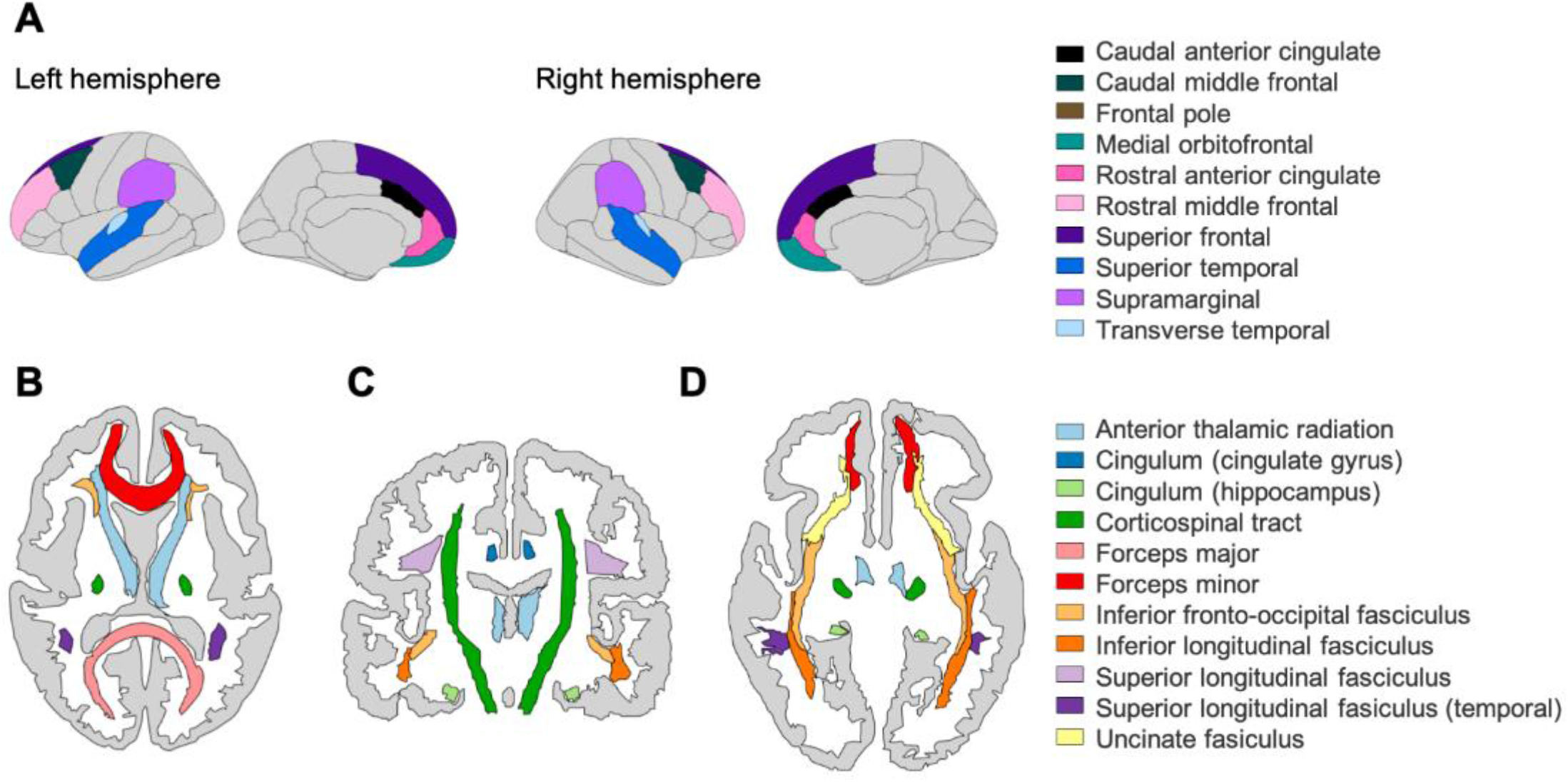
Top: A) Grey matter ROIs based on the DK atlas (cortical volume, N=246) in the left and right hemisphere. Bottom: White matter ROIs based on the John’s Hopkin’s University atlas (fractional anisotropy, N=165) in B) Transverse plane (superior), C) Coronal plane and D) Transverse plane (inferior).

### Network Estimation Methods

All statistical analyses and plots were completed using R (R Core Team 2020) version 3.6.3 (“Holding the Windsock”). Network estimation was performed using the packages bootnet (version 1.4.3, Epskamp and Fried 2020), igraph (version 1.2.6, Amestoy et al. 2020), qgraph (version 1.6.5, Epskamp et al. 2020), and networktools (version 1.2.3, Jones 2020). We used these tools to estimate weighted partial correlation networks, which allowed determination of conditional dependencies among our cognitive and neural variables. For example, in a multilayer network, any partial correlation between node A (e.g., matrix reasoning) and node B (e.g., the caudal anterior cingulate) is one that remains *after* controlling for the associations among A and B with every other node in the network (e.g., other cognitive abilities and cortical volume ROIs). To estimate these networks, we applied Gaussian Graphical Models (Pearson correlations) using regularization (graphical lasso, see Friedman, Hastie, and Tibshirani 2008) with a threshold tuning parameter of 0.5 and pairwise deletion to account for missingness. These methods have been widely used to generate sparser networks by penalizing for more complex models—thus, decreasing the risk of potentially spurious (e.g., false positive) connections and enabling simpler visualization and interpretation of conditional dependencies between nodes (Epskamp and Fried 2018). We hypothesized that our results would show positive partial correlations (in line with mutualism theory) both within cognitive (e.g., as observed in Mareva and Holmes 2020 and Schmank et al. 2019) and within neural measures (single-layer networks) as well as between brain-behavior variables in the multilayer networks.

Note that age was included as a node in the estimation procedures of all partial correlation networks (i.e., edge weights, centrality, network stability, and community detection) but was not included in the visualizations of our networks and centrality plots, or in network descriptive statistics (i.e., mean, median, and range of edge weights). For a comparison of the use of age (i.e., included in estimation or regressed out beforehand), see the Supplementary Material.

### Node Strength Centrality (Single-layer Networks)

To assess the statistical interconnectedness or connectivity of cognitive and neural nodes relative to their neighbors within our single-layer networks, we estimated node strength, a weighted degree centrality measure calculated by summing the absolute partial correlation coefficients (edge weights) between a node and all other nodes it connects to within the network. Note that our brain structural covariance networks involve ROIs that are not necessarily anatomically connected, preventing certain inferences such as information flow. Nodes were classified as central if the magnitude of their strength z-score was positive and equal to or greater than one standard deviation above the mean. We do not discuss or interpret negative centrality z-score values for our single-layer networks.

### Community Detection and Bridge Strength Centrality (Multilayer Networks)

In our multilayer networks, we applied the Walktrap community detection algorithm (Pons and Latapy 2005) to determine in a data-driven manner whether clustering, or grouping, of nodes (e.g., cognitive and/or neural) occurred. The Walktrap algorithm assesses how strongly related nodes are to each other (that can be due to similarity, e.g., because nodes A and B are similar, or it can be because nodes A and B are different but node A has a strong impact on node B; see “*Topological overlap and missing nodes*” of Fried and Cramer 2017). The Walktrap algorithm works by taking recursive random walks between node pairs and classifies communities according to how densely connected these parts are within the network (wherever the random walks become ‘trapped’). Walktrap is widely used in the network psychometrics literature and, in a Monte Carlo simulation study, was shown to outperform other algorithms (e.g., InfoMap) for sparse count networks (e.g., those used in diffusion tensor imaging), although it must be noted that this result was for networks made up of 500 nodes or higher (Gates et al. 2016). We also calculated the maximum modularity index value (Q), which estimates the robustness of the community partition (Newman 2006). We interpreted values of 0.5 or above as evidence for reliable grouping.

Instead of traditional absolute strength, we calculated bridge strength, a novel weighted degree centrality measure originality developed to study comorbidity between mental disorders (see Jones, Ma, and McNally 2019 for overview). Bridge strength centrality sums the absolute value of every edge that connects one node (e.g., matrix reasoning) in one pre-assigned community (e.g., cognition) to another node (e.g., caudal anterior cingulate) in another pre-assigned community (e.g., brain). Recent simulation work has shown that the method can reliably recover true structures of bridge nodes in both directed and undirected networks (Jones, Ma, and McNally 2019). Rather than relying on straightforward ‘brain’ or ‘behavior’ assignments to classify nodes, we pre-assigned communities for bridge strength calculation based on results from the Walktrap algorithm.

The presence of bridges between communities (e.g., if nodes from topological distinct clusters such as cognition vs. brain feature relations) might suggests the existence of intermediate endophenotypes (Fornito and Bullmore 2012; Kievit et al. 2016), and potentially identify potential nodes (both cognitive and neural) that might one day guide intervention studies. Nodes were classified as central if the magnitude of their strength z-score was positive and equal to or greater than one standard deviation above the mean. We do not discuss or interpret negative centrality z-score values for our multilayer networks.

### Node Centrality Stability (Single and Multilayer Networks)

Lastly, we quantified the reliability of our centrality estimates for all single-layer (absolute strength of cognitive and brain structural covariance nodes) and multilayer networks (bridge strength). We estimated the correlation stability (CS) coefficient, calculated as the maximum proportion (out of N=2,000 bootstraps) of the sample that can be dropped out and, with 95% probability, still retain a correlation of 0.7 (correlation between rank order of centrality in network estimated on full sample with order of subsampled network in smaller N), with a CS value of 0.5 considered to be stable (Epskamp, Borsboom, and Fried 2018). Lastly, also using bootstrapping, we determined the stability of the edge-weight coefficients but present these results in the Supplementary Material.

## RESULTS

### Single-layer Network Models (Cognitive, Cortical Volume and Fractional Anisotropy)

The regularized partial correlation (PC) network for the CALM cognitive data is shown in Figure 4 (top left). This network shows that all partial correlations are positive, and most have small magnitude (mean PC=0.08, median PC=0.07, PC range=0—0.63). One edge (between reading and spelling) was an outlier (PC=0.63, all others are between 0 and 0.27), likely due to close content overlap (verbal ability). Regarding centrality, three nodes emerged as strong (positive z-score at or greater than one standard deviation above the mean): (in descending order of centrality strength) reading, numerical operations, and peabody picture vocabulary test (Figure 4 top right). Overall, centrality estimates were stable, indicated by a high correlation stability (CS)-coefficient of 0.75, revealing that at least 75% of the sample could be dropped while maintaining a correlation of 0.7 with the original sample at 95% probability.

**Figure 4.**
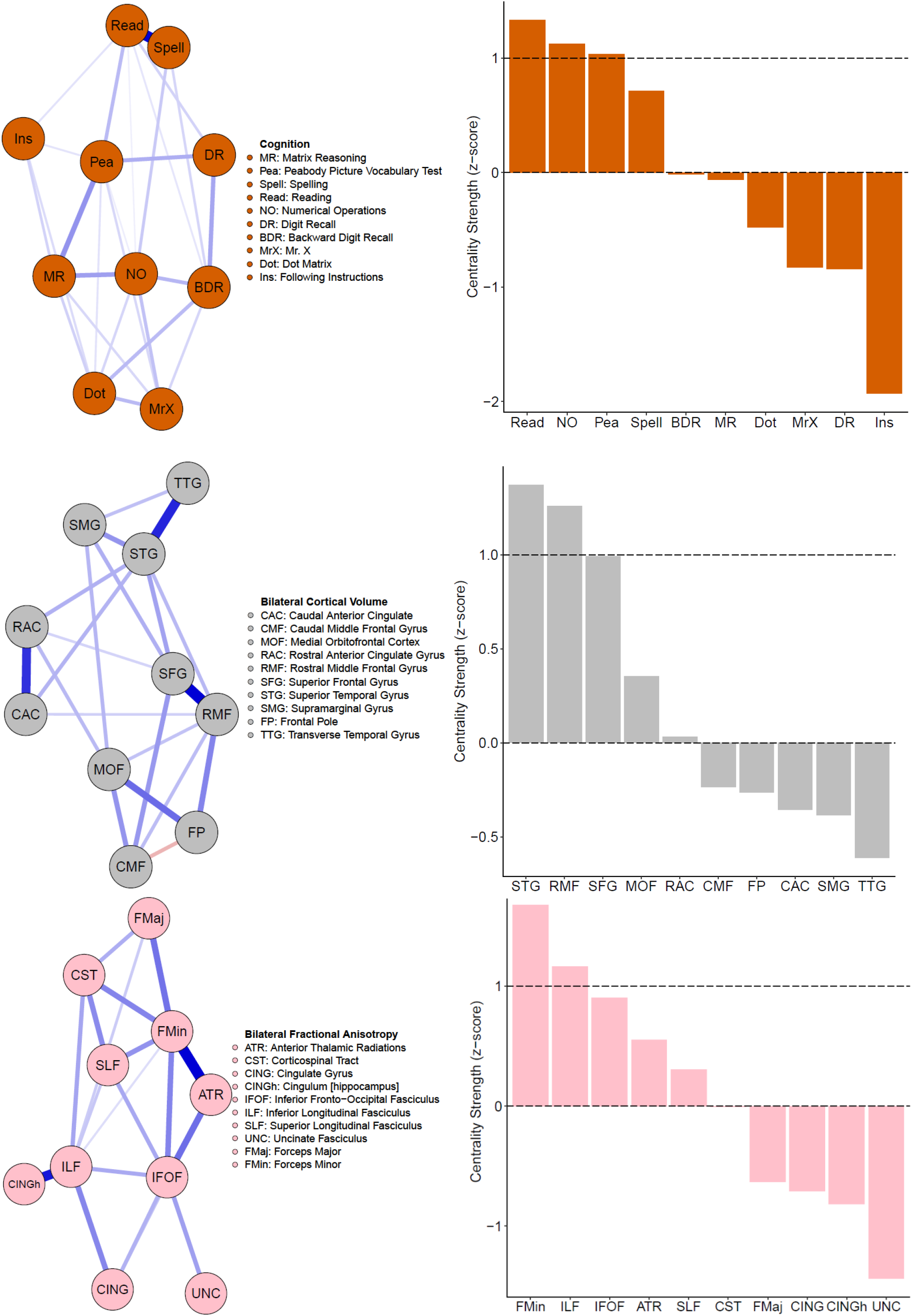
Single-layer partial correlation networks. Top: Network visualization (spring layout, left side) of CALM cognitive data (N=805). Centrality estimates (z-scores) of all cognitive tasks (right). Middle: Network visualization (spring layout, left side) of CALM cortical volume data (N=246). Centrality estimates (z-scores) of all cortical volume nodes (right). Bottom: Network visualization (spring layout, left side) of CALM fractional anisotropy data (N=165). Centrality estimates (z-scores) of all fractional anisotropy nodes (right).

Next, we estimated the partial correlation network among 10 grey matter regions as shown in Figure 3 (top) above. All edges weights (mean PC=0.09, median PC=0, PC range= −0.15—0.52) of the cortical volume network (Figure 4, middle left) were positive apart from one negative path (caudal middle frontal gyrus and frontal pole PC= −0.15). Note, the negative path between the caudal middle frontal gyrus and frontal pole might be due to the frontal pole correlating surprisingly weakly with other grey matter nodes and displaying a steeper decrease pattern across age (Figures 1 and 2). Two ROIs emerged as central (in descending order of centrality strength): superior temporal gyrus and rostral middle frontal gyrus (Figure 4, middle right). Similar to the cognitive network, cortical volume centrality was stable (CS-coefficient=0.52), indicating that about 52% of the sample could be subtracted to maintain a correlation of centrality estimates above 0.7 compared to the full sample. This finding is despite the lower sample size compared to the behavioral data (N=805 for behavior vs. N=246 for cortical volume).

Finally, similar to the cognitive and the grey matter covariance network, the fractional anisotropy network (Figure 4, bottom left) has positive partial correlations with all edge weights varying between small and moderate values: mean PC=0.08, median PC=0, and PC range=0—0.44. Two white matter ROIs displayed centrality (Figure 4, bottom right). These included (in descending order) the forceps minor and inferior longitudinal fasciculus. Finally, fractional anisotropy centrality was moderately stable (CS-coefficient=0.44) indicating that about 44% of the sample could be removed while maintaining a 0.7 association with 95% probability. This is possibly due to the much lower sample size (N=165) compared to the cognitive (N=805) and grey matter (N=246) networks.

### Bridging the Gap: Multilayer Networks

The regularized partial correlation network analyses for the CALM multilayer networks data are shown in Figure 5. Consistent with the pattern found in the single-layer networks, the cognitive and grey matter multilayer network (top left of Figure 5) edges are mostly positive and small to moderate weights (mean PC= 0.04, median PC=0, PC range= −0.12—0.64). Comparably, the cognitive and white matter multilayer network (Figure 5, top right) had similar edge weight estimates (mean PC=0.04, median=0, range= −0.2—0.65). Finally, combining all measures together (tri–layer network consisting of cognition, grey and white matter, bottom center of Figure 5) produced a network with similar characteristics to the bi-layer networks (mean PC=0.02, median PC=0, PC range= −0.2—0.66). For the bi-layer networks, the Walktrap algorithm produced either three (cognition-white matter) or four (cog-grey matter) clusters that consisted entirely of cognitive or neural nodes except for following instructions (Ins), which was either kicked out (cognition-grey matter, Q=0.56, indicating strong modularity) or grouped with a neural node (forceps minor of the cognition-white matter, Q=0.39, indicating moderate modularity). The result for the tri-layer network (Q=0.25, indicating weak modularity) was more complex with a total of 15 communities (Figure 5, bottom center; note age was found to be in a community by itself but is not shown in the figure).

**Figure 5.**
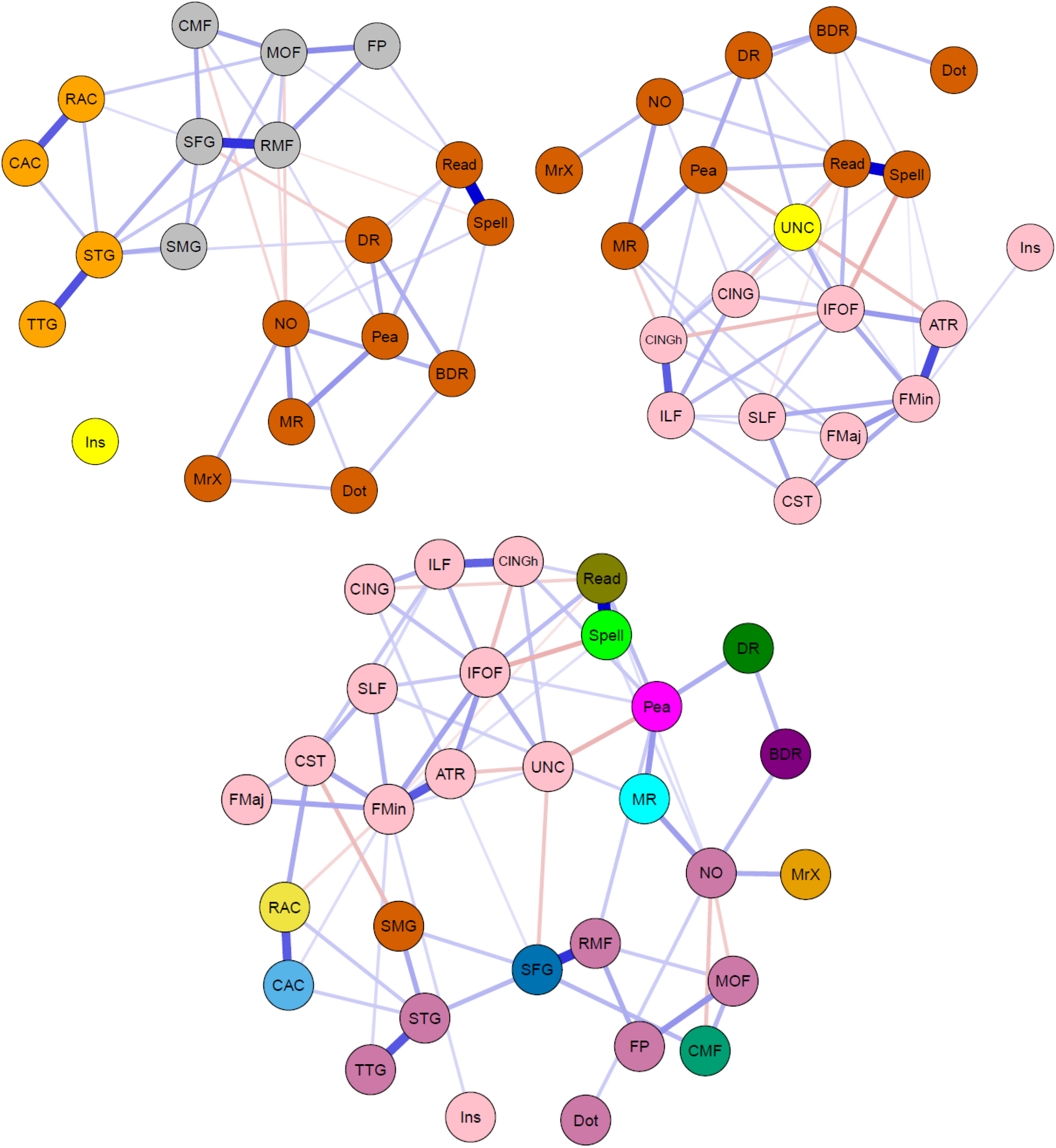
Network visualizations (spring layout) of partial correlation multilayer networks for CALM data. Colors indicate groups determined by the Walktrap algorithm (see above). Top: Bi-layer networks consisting of cognition and grey matter (top left), and cognition and white matter (top right). Bottom: Tri-layer network consisting of cognition, grey matter and white matter (center).

Regarding centrality, we report bridge strength (Figure 6). In the cognitive-grey matter network, three bridge nodes surfaced (in descending order: superior temporal gyrus, superior frontal gyrus, and rostral middle frontal gyrus, Figure 6 top left). In terms of stability, the CS-coefficient was 0.20, indicating that the bridge strength estimates were unstable under bootstrapping conditions. In the cognitive-white matter bi-layer network, three nodes (in descending order: uncinate fasciculus, inferior frontal-occipital fasciculus, and hippocampal cingulum) emerged as possible bridge nodes (Figure 6, top right). Moreover, the centrality estimates had a CS-coefficient of 0.13, once again suggesting that the bridge strength estimates were unstable. Lastly, for the tri-layer network, five nodes displayed positive bridge strength equal to or greater than one standard deviation above the mean (Figure 6, bottom center). These included (in descending order): reading, peabody picture vocabulary test, superior frontal gyrus, spelling, and numerical operations. Much better than the bi-layer networks, the tri-layer network bridge strength estimates were moderately stable (CS-coefficient=0.44).

**Figure 6.**
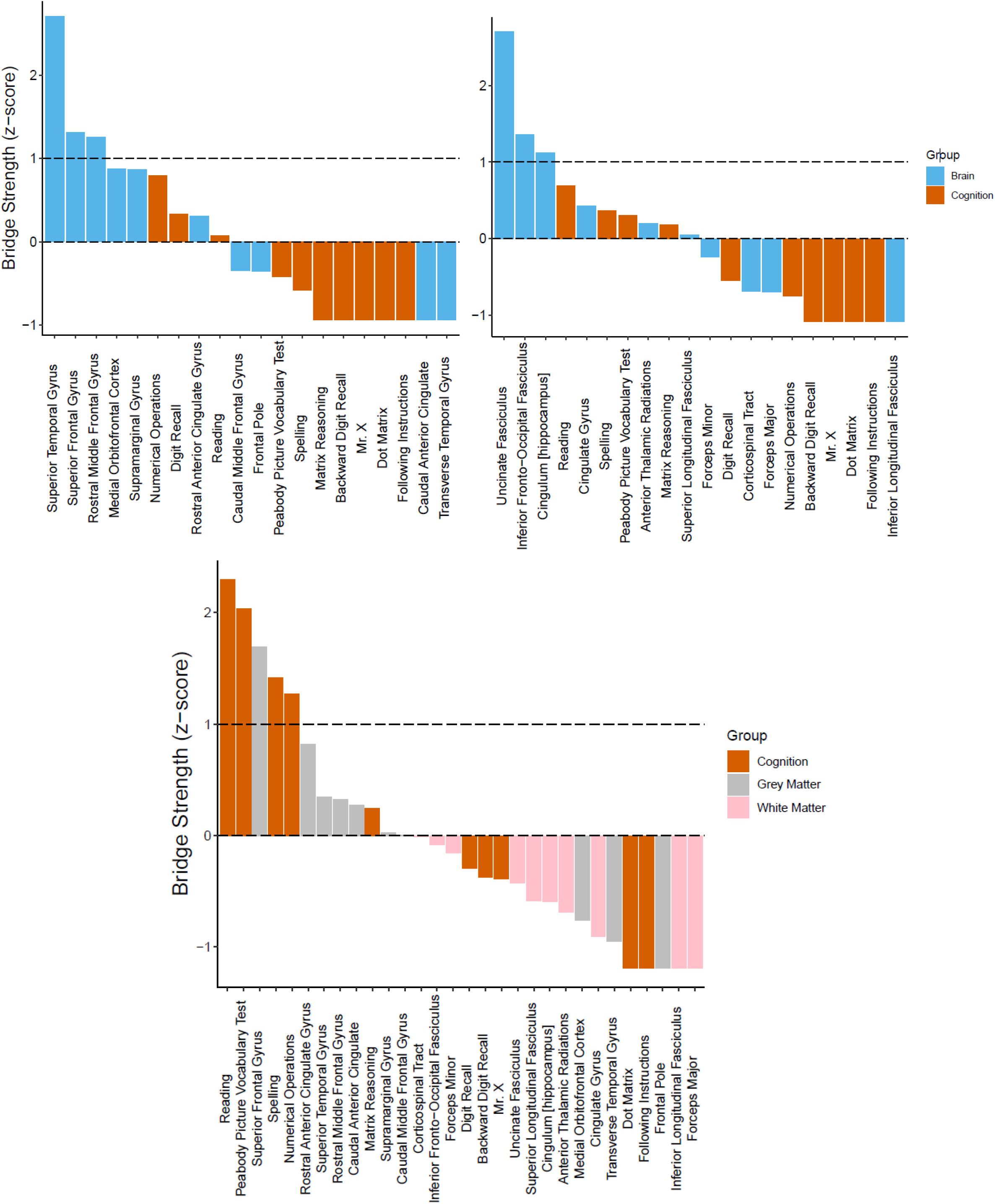
Bridge centrality estimates (z-scores) for multilayer networks. Top: Bi-layer networks consisting of cognition and grey matter (top left), and cognition and white matter (top right). Bottom: Tri-layer network consisting of cognition, grey matter and white matter (center). Dashed lines indicate mean strength and one standard deviation above the mean.

## DISCUSSION

### Summary of Main Findings

In this study, we used network analysis (partial correlations) to examine the neurocognitive structure of general intelligence in a childhood and adolescent cohort of struggling learners (CALM). For our single-layer networks (Figure 4), we found that cognitive, grey matter and white matter networks contained mostly (if not all) positive partial correlations. Moreover, in all single-layer networks, at least two nodes emerged as more central than others (as indexed by node strength equal to or greater than one standard deviation above the mean), which varied in stability from moderately to highly reliable. In the cognitive network, this included a verbal ability (specifically reading and peabody picture vocabulary test) and crystallized intelligence (i.e., numerical operations). In the structural brain networks (grey matter cortical volume and white matter fractional anisotropy), two nodes emerged as central for the grey matter network (superior temporal gyrus and rostral middle frontal gyrus) and white matter network (forceps minor and inferior longitudinal fasciculus) passed the centrality threshold. Furthermore, we extended previous approaches by integrating networks of structural brain data with a cognitive network, forming bi- and tri-layer networks (Figure 5). Doing so, we observed multiple (both positive and negative) partial correlations between brain and behavior variables. Using bridge strength as a metric, we found that, in our bi-layer networks, only neural nodes harbor significant connections across communities (defined by the Walktrap algorithm) and levels of organization (Figure 6, top). In contrast, in the tri-layer network, we found support that mostly cognitive nodes connect across different communities (Figure 6, bottom). Overall, our results suggest which behavioral and neural variables have greater (possible) influence among or might be more influenced by other nodes and potentially serve as bridges between the brain and cognition within general intelligence. However, the literature on drawing inferences from networks to the most likely consequences of intervening on the network is complex and rapidly changing, (e.g., Dablander and Hinne 2019; Henry, Robinaugh, and Fried 2020; Levine and Leucht 2016).

### Interpretation of Network Models and Community Detection Analyses

For the cognitive network, each node corresponded to a single cognitive task (e.g., matrix reasoning) while partial correlations (weighted edges) between nodes were interpreted as compatible with (possible) causal consequences of interactions among cognitive abilities during development. This interpretation is compatible with the mutualism theory of cognitive development (van der Maas et al. 2006), whereby cognitive abilities positively reinforce each other (e.g., positive partial correlations) over time to produce the positive manifold (Spearman 1904). Mutualism hypothesizes that general intelligence emerges from causal interactions among abilities rather than a general latent factor (Fried 2020b; Kan, van der Maas, and Levine 2019). Hence, cognition is viewed as a complex system derived from the dynamic relations of specific abilities that become more intertwined over development.

The existence of only positive edges in our cognitive network would be expected under a mutualistic perspective (interactions among cognitive variables), which at its essence is a network theory of general intelligence, although longitudinal analyses are needed to further substantiate this claim. Initially, it was surprising that two of the three most central nodes (i.e., reading and peabody picture vocabulary test) relate to verbal ability rather than abilities such as fluid intelligence and working memory (matrix reasoning and (forward and backward) digit recall), which are traditionally viewed as causal influences on cognitive development (Cattell 1971). However, an emerging body of literature suggests that verbal ability plays a crucial role in cognitive development (e.g., between reading and working memory before 4^th^ grade, Peng et al. 2018 and Zhang and Malatesha Joshi 2020, as well as driving the emergence of reasoning (Kievit, Hofman, and Nation 2019; also see Gathercole et al. 1999)).

As for our neural networks (here, grey matter cortical volume and white matter fractional anisotropy), individual nodes were comprised of a single ROI. Importantly, we did not interpret weighted edges as an index of direct connectivity. Instead, the presence of strong associations between these ROIs would be compatible with the hypothesis of coordinated development (see Alexander-Bloch, Giedd, and Bullmore 2013) whereby certain brain regions show preferential correlations to each other than more peripheral regions over time (e.g., childhood to late adolescence) as well as the notion of “rich” (Heuvel and Sporns 2011) and “diverse” (Bertolero, Yeo, and D’Esposito 2017) clubs that enable local and global integration, respectively. The most central node, the superior temporal gyrus which has been implicated in verbal reasoning (e.g., Khundrakpam et al. 2017). Regarding white matter, the two strongest nodes (forceps minor and inferior longitudinal fasciculus) while not anatomically close, instead represent long-range connections (see de Mooij et al. 2018) that have been linked to mathematical ability (Navas-Sánchez et al. 2014) and visuospatial working memory (Krogsrud et al. 2018).

Finally, we integrated both domains (cognitive abilities and brain metrics) into combined multilayer networks (cognition-grey matter, cognition-white matter and cognition-grey and- white matter). Doing so allowed us to attempt comparison and integration simultaneously across explanatory levels within the same analytical paradigm (network analysis) and statistical metrics (partial correlations, centrality, and community detection). From this analysis, three observations immediately stood out. First, there were multiple partial correlations between cognitive and neural nodes (especially in the cognitive-white matter and cognitive-grey matter and- white matter networks). Second, compared to the single-layer networks, the multilayer networks have more negative partial correlations. Together, these two findings further suggest that associations between the brain and cognition are complex as they defy straightforward (e.g., only positive and/or one-to-one) relationships and interpretations. However, it should be noted that causality (e.g., conditioning on colliders, see Rohrer 2018 for overview of interpretations of correlations in graphical causal models in observational data) becomes even more difficult to determine with networks incorporating multiple levels of organization (e.g., cognition and structural brain covariance). Finally, we found a peculiar role of the cognitive task following instructions (Ins) within all multilayer networks. For example, in the cognitive-grey matter network, Ins had no partial correlations with any other nodes within the network while in both the cognitive-white matter and tri-layer network (cognition, grey and white matter) Ins *only* correlated with the forceps minor (FMin), a neural node, and not any of the cognitive variables. This might suggest that following instructions, traditionally a working memory task and often analyzed using structural equation modeling, may have distinct psychometric properties (e.g., one-to-one mapping) when compared to other cognitive tasks when modeled through network science approaches, and/or when adjusted for all shared correlations.

Further inspection of bridge strength centrality showed an interesting pattern: (discounting the one standard deviation cutoff) the neural nodes are stronger than the cognitive variables within the multilayer networks, despite there being an equal number of cognitive nodes for each brain metric. This is possibly due to the large number of edges between them (grey and white matter regions) and both cognitive and other neural nodes. In other words, since the neural nodes contain a larger number of connections (partial correlations) across explanatory levels, they display greater bridge strength (bridge strength sums inter-network correlations).

In other ways, the multilayer networks differed. First, in the tri-layer network, the four of the five central nodes were cognitive variables while, in the bi-layer networks, the central nodes were neural ROIs. Three of these central cognitive nodes in the tri-layer network (reading, peabody picture vocabulary test, and numerical operations) were also found to be central in the single-layer cognitive network. This further suggests the importance of mathematical and verbal ability in understanding the cognitive neuroscience of general intelligence. Secondly, the fact that cognitive nodes were found to be central only in the tri-layer network suggests that grey and white matter, while related, possibly reveal unique information about cognition when combined and analyzed together simultaneously.

### Limitations of the Current Study

This study contains several limitations that require caution when interpreting the results. First and foremost, these findings are based on cross-sectional data. While adequate to help tease apart individual differences in cognition *between* people, cross-sectional data cannot be used to elucidate differences in *changes within* individuals over time, such as during development. Therefore, longitudinal analyses are needed before attempting to make strong inferences about the dynamics of these networks. Reiterating this point, a recent study using intelligence data (Schmiedek et al. 2020) found that a cross-sectional analysis of the *g* factor of cognitive ability was unable to capture within-person changes in cognitive abilities over time. This finding further stresses the necessity to integrate cross-sectional (between-person) differences and longitudinal (within-person) changes when studying cognition.

Moreover, the CALM sample represents an atypical sample (Holmes et al. 2019), with participants who consistently score lower on measures of attention, learning and/or memory than age-matched controls (see Figure 2 (Level I) of Simpson-Kent et al. 2020 for comparison to a typically developing sample). As a result, these analyses would need to be replicated in additional (ideally larger) samples with different cognitive profiles before our results can be generalized. This shortcoming of the present study is echoed by the low stability estimates found for the centrality values in the bi-layer networks, which might be due to the differences between the sample sizes of the neural data (grey matter, N=246; white matter, N=165) compared to cognition (N=805). Interestingly, the tri-layer network showed moderate bridge strength stability, but also displayed weak modularity. Moreover, given that the Walktrap algorithm produced 15 communities in the network, which contained only 31 nodes (including age), we further state that this result should be interpreted with caution and must be corroborated in larger cohorts (e.g., ABCD study, Casey et al. 2018).

Lastly, we re-ran our analyses to test the sensitivity of our main findings (e.g., positive partial correlations and central nodes) to potential outliers (defined as ± 4 standard deviations). Doing so did not severely alter the partial correlation weights between nodes in our networks (see Supplementary Material for detailed comparisons). It must be restated that our data comes from an atypical sample, which might influence brain metrics even with rigorous quality control procedures. Therefore, despite this discrepancy, our data supports brain-behavior ‘bridges’ in general intelligence.

### Future Directions Toward Theory Building in Cognitive Neuroscience

Our results that suggest verbal abilities rather than fluid intelligence or working memory might play a more pivotal role in the development of cognitive ability fits with the gradual progression in schooling. For example, before children can successfully be taught more advanced subjects (e.g., history, reading comprehension, etc.), they must first become competent in basic language faculties. In other words, it may be that verbal skills (e.g., reading and spelling) facilitate performance on abstract tests, even in the absence of direct knowledge-based task demands. Recent evidence has been found supporting this notion and suggest that verbal ability, particularly reading and vocabulary in relation to working memory and reasoning, might drive early cognitive development (Kievit, Hofman, and Nation 2019; Peng et al. 2018; Zhang and Malatesha Joshi 2020). Therefore, future studies could further examine whether greater verbal ability in early development facilitates greater acquisition of higher-level cognitive skills by lowering computational demands in working memory.

Moreover, in this context, the fact that the numerical operations task was also found to be central (tri-layer network only) should be expected since mathematics (e.g., arithmetic) also involves symbol manipulation. In terms of mutualism (van der Maas et al. 2006), future models (ideally in longitudinal samples) could test whether language and other symbolic abilities show progressively higher reciprocal associations during early development compared to other abilities until more complex cognition (i.e., fluid reasoning and working memory) develops in later childhood (also see Kievit, Hofman, and Nation 2019 and Peng et al. 2018).

We argue that future studies should aim to incorporate data from different scales, not only temporal (e.g., development) but also levels of organization (e.g., brain and behavior). Furthermore, results from different levels can more easily be interpreted if these datasets are analyzed using a unified quantitative framework that combines strengths from various statistical techniques (such as pairwise and partial correlations to reveal causality in brain functional connectivity networks, see Reid et al. 2019). Last, and perhaps most important, cognitive neuroscientists must formulate mechanistic (e.g., Bertolero et al. 2018) and generative models (for instance, Akarca et al. 2020) to gain further insights from past and help guide future controlled experiments. Researchers must not shy away from but rather embrace the complexity of the brain and cognition (see Fried and Robinaugh 2020 for similar argument for mental health research). Intelligence is a complex system—to understand it, we must treat it as such.

## ACKNOWLEDGMENTS

The Centre for Attention Learning and Memory (CALM) research clinic is based at and supported by funding from the MRC Cognition and Brain Sciences Unit, University of Cambridge. The Principal Investigators are Joni Holmes (Head of CALM), Susan Gathercole (Chair of CALM Management Committee), Duncan Astle, Tom Manly, Kate Baker and Rogier Kievit. Data collection is assisted by a team of researchers and PhD students at the CBU that includes Joe Bathelt, Giacomo Bignardi, Sarah Bishop, Erica Bottacin, Lara Bridge, Annie Bryant, Sally Butterfield, Elizabeth Byrne, Gemma Crickmore, Edwin Dalmaijer, Fanchea Daily, Tina Emery, Laura Forde, Grace Franckel, Delia Fuhrmann, Andrew Gadie, Sara Gharooni, Jacalyn Guy, Erin Hawkins, Agniezska Jaroslawska, Amy Johnson, Jonathon Jones, Silvana Mareva, Elise Ng-Cordell, Sinead O’Brien, Cliodhna O’Leary, Joseph Rennie, Ivan Simpson-Kent, Roma Siugzdaite, Tess Smith, Stepheni Uh, Francesca Woolgar and Mengya Zhang. The authors wish to thank the many professionals working in children’s services in the South-East and East of England for their support, and to the children and their families for giving up their time to visit the clinic.

I.L.S.-K. is supported by the Cambridge Trust., E.I.F. did not receive funding for this project, S.M. by the Medical Research Council PhD Studentship and D.A. by the Medical Research Council Doctoral Training Partnership Studentship (Project Code: SUAH/012; Award: RG86932) and the Cambridge Trust. E.T.B. is supported by an NIHR Senior Investigator award. This project also received funding from the European Union’s Horizon 2020 research and innovation programme (grant agreement number 732592). R.A.K. is supported by the Wellcome Trust (Grant No. 107392/Z/15/Z), the UK Medical Research Council SUAG/014 RG91365 and the Hypatia Fellowship by Radboud University.

## Citation Diversity Statement

Recent work has demonstrated that gender^1,2^ (as well as racial/ethnic^3^) minorities are systemically under-cited in neuroscience. We estimated the gender citation practices of this manuscript using the *Journal of Cognitive Neuroscience*’s Gender Citation Balance Index tool (GCBI-alyzer, https://postlab.psych.wisc.edu/gcbialyzer/). Overall, 56 DOIs were successfully categorized and metrics for man (first author)/man (last author), woman (first author)/man (last author), man (first author)/woman (last author), and woman (first author)/woman (last author) were calculated. Percentages for the 56 DOIs were as follows: MM=58.9%, WM=17.9%, MW=12.5%, and WW=10.7%. The GCBIs were as follows: MM=0.444, WM= −0.467, MW=0.157, and WW= −0.281. This indicates that we over-cited MM and MW (>0 indicates over-citation) and under-cited WM and WW (<0 denotes under-citation). Note that these estimates assume a binary paradigm of gender (man or woman) and, therefore, do not account for non-binary identifications of gender. Furthermore, these estimates are based on the authorship practices of one journal, *Journal of Cognitive Neuroscience*. Even though this manuscript can be classified as (developmental) cognitive neuroscience, citations also include purely behavioral (e.g., cognitive psychology) and neuroscience (e.g., physical properties of the brain such as small-worldness) literature.

Nevertheless, the estimated metrics indicate that we have under-cited literature involving women and disproportionately over-cited research involving men. Lastly, although we will not add references merely to ensure a desirable proportion, we will thoroughly examine any overlooked yet pertinent literature during the resubmission/revision phase, including expanding our literature search to a wider net of journals, to seek out relevant literature.

## SUPPLEMENTARY MATERIAL

### Edge-weight Stability Analyses

To further quantify the reliability of our partial correlation network edge-weights, we performed bootstraps (N=2,000) and compared the bootstrapped mean values to the original sample estimates (Supplementary Figures 1-3). We do not show the bootstraps for the multilayer networks due to the size of the plots but they (and all code for this project) can be found online (https://osf.io/36d2n/). Bootstrapped edge-weight means were consistently near the original sample value with the most variable being the white matter network (Supplementary Figure 3) and the multilayer networks (not shown). The low edge-weight stability in these networks could possibly due to lower sample sizes of neural data (especially in the white matter network, N=165, although centrality strength was moderately stable, CS-coefficient=0.44), including when structural brain and cognitive data were combined. This, in turn, could have influenced the low stability estimates of the bridge centrality values in the multilayer networks.

**Supplementary Figure 1.**
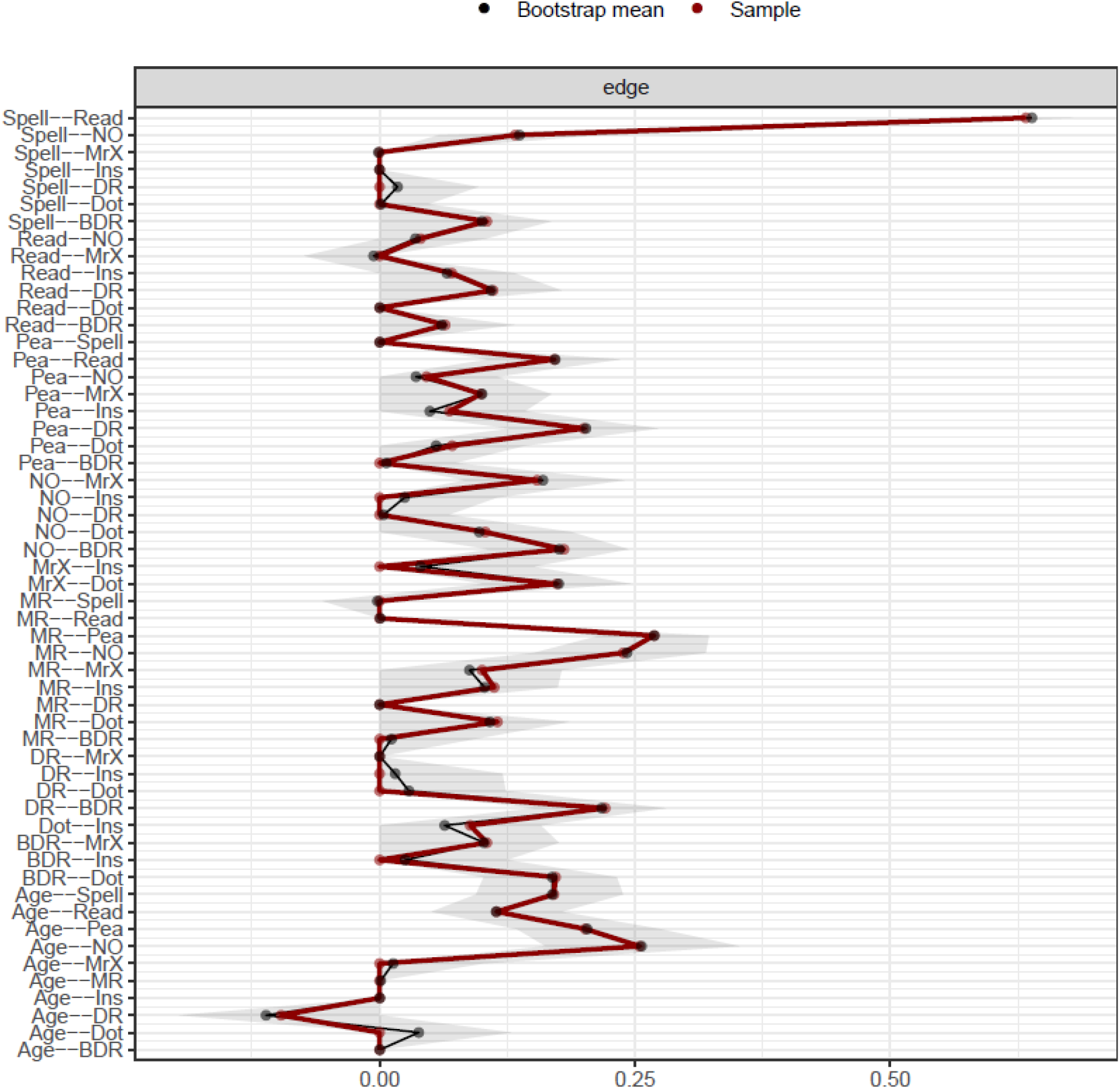
Comparisons between bootstrapped means and original sample edge-weight estimates for the CALM cognitive partial correlation network.

**Supplementary Figure 2.**
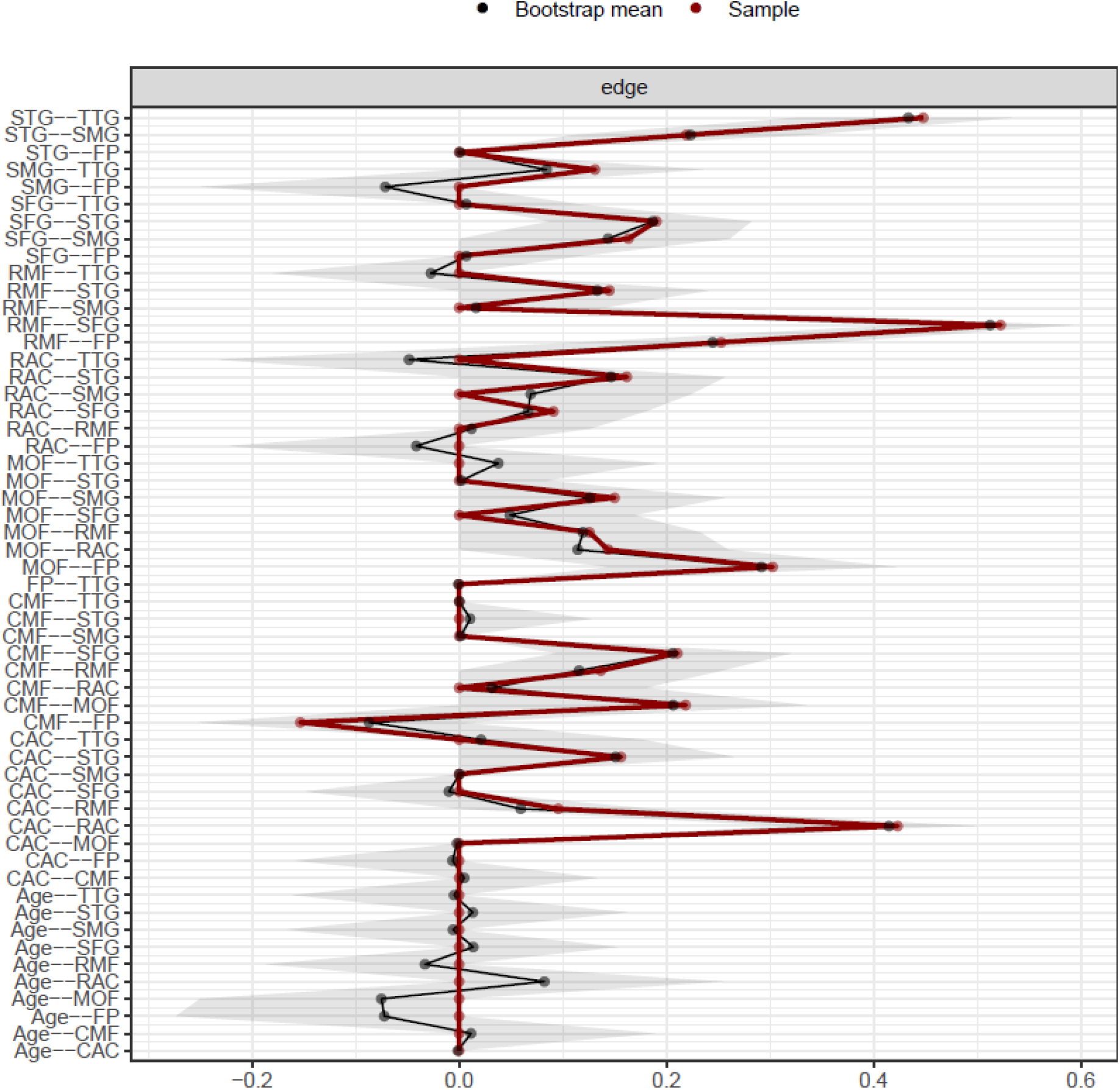
Comparisons between bootstrapped means and original sample edge-weight estimates for the CALM grey matter partial correlation network.

**Supplementary Figure 3.**
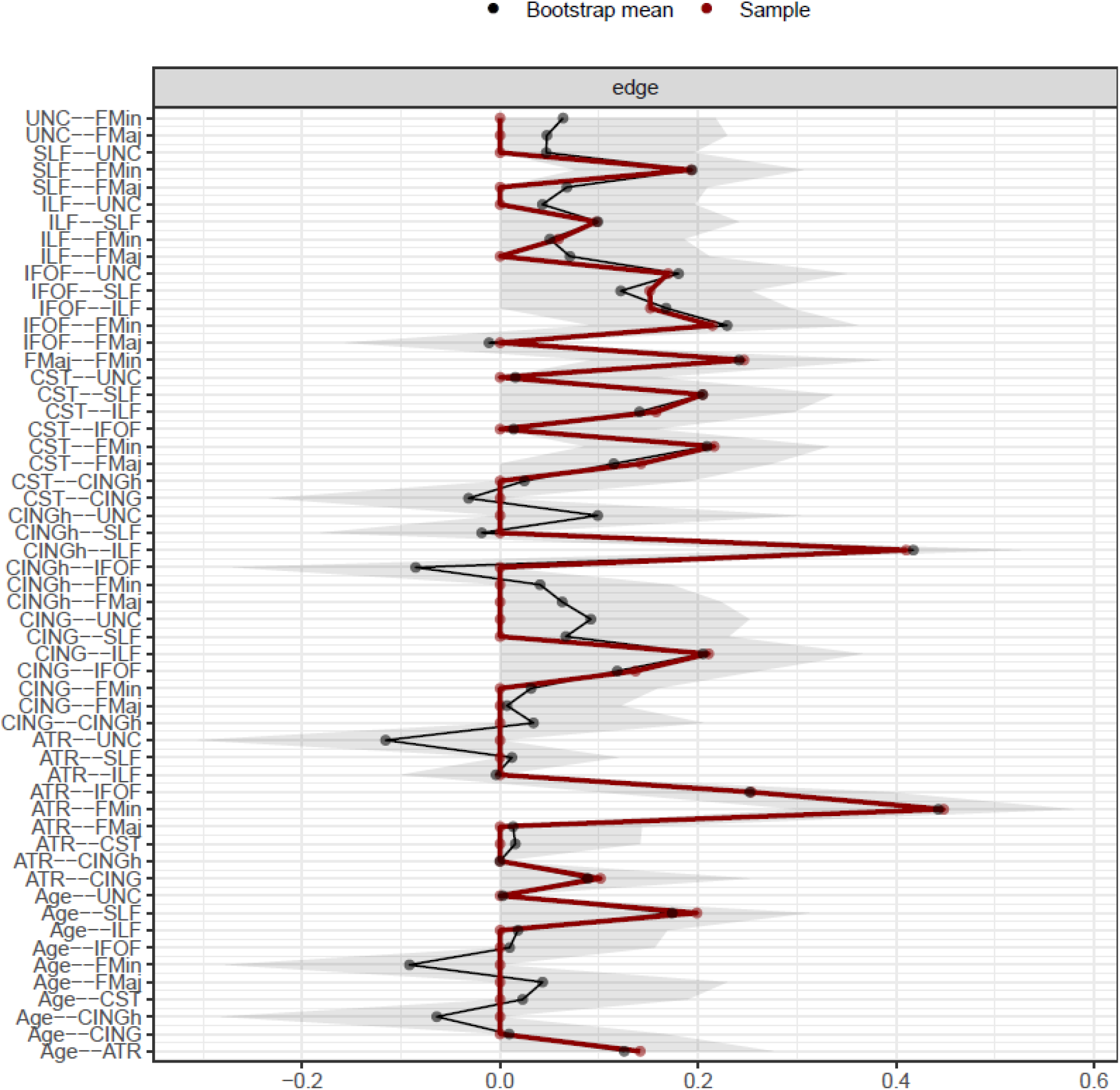
Comparisons between bootstrapped means and original sample edge-weight estimates for the CALM white matter partial correlation network.

### The Possible Effect of Outliers on Major Findings

In a <u>previous version</u> of this manuscript, we observed that two FA values (1 for the uncinate fasciculus, 1 for the forceps major), which represent potential outliers with undue influence on the partitioning of the Walktrap algorithm in the single-layer white matter network. Removing this data yielded a distinct, and more parsimonious clustering solution (2 communities vs. 5). Moreover, removing this outlier did not affect any summary statistics for the white matter partial correlation (single-layer) network except for range. Nevertheless, below we present the Pearson correlations between the weights obtained from the original data presented in the main manuscript and those from the data after all outliers (defined as ± 4 standard deviations) are removed (Supplementary Table 1). Due to the vast similarity in descriptive statistics and high correlations between partial correlation weights, we conclude that outliers did not confound the results of this study. However, it must be noted that outliers might slightly affect community detection, but we chose to keep the original data due to the nature of our sample (struggling learners, therefore behavioral and neural data might be atypical to begin with) and given the fact that the neural data was already quality controlled. Furthermore, the two outlier white matter ROIs occurred in two separate particpants (1 outlier each) while the rest of their ROIs were consistent with the rest of the sample. In close, we argue that outliers (both cognitive and neural) are likely not due to measurement error but instead represent realistic values of an atypically developing sample.

**Supplementary Table 1.** Comparisons between partial correlation (PC) networks (original data vs. outliers removed). These include summary statistics such as mean, (standard deviation), [range] and Pearson correlations between PC graph weights using pairwise complete observations to account for missingness.

| **Network Type** | **Original Data** | **Outliers Removed** | **Pearson Correlation** |
| --- | --- | --- | --- |
| Cognitive | 0.08 (0.11)<br>[0, 0.63] | 0.08 (0.11)<br>[0, 0.61] | 0.99 |
| Grey Matter | 0.09 (0.14)<br>[-0.15, 0.52] | 0.09 (0.14)<br>[-0.15, 0.52] | 1 |
| White Matter | 0.08 (0.11)<br>[0, 0.44] | 0.08 (0.13)<br>[-0.14, 0.47] | 0.93 |
| Cognitive-grey matter | 0.04 (0.1)<br>[-0.12, 0.64] | 0.03 (0.09)<br>[-0.11, 0.62] | 0.97 |
| Cognitive-white matter | 0.04 (0.1)<br>[-0.2, 0.65] | 0.04 (0.1)<br>[-0.22, 0.65] | 0.97 |
| Tri-layer | 0.02 (0.08)<br>[-0.2, 0.66] | 0.02 (0.08)<br>[-0.19, 0.65] | 0.98 |

### How to Deal with Age?

As in previous literature, in our sample age shows a clear positive association with intelligence measures and brain structure (Figure 2). This fact, however, may further complicate any interpretations of (possible) causal interactions between cognitive and/or neural nodes. This leaves us with at least two options of how to deal with the relationship of age to cognitive ability, and grey and white matter structural covariance: 1) We could estimate the partial correlation network and include age as a node, therefore, choosing to estimate it *simultaneously* with the cognitive and neural variables (this is the option we chose for the main manuscript analyses), or 2). We could *regress out* the association of age for each variable (age would show no correlation for cognitive and/or neural measures) *before* network estimation. Both approaches have corresponding pros and cons. For instance, choosing to include age has the benefit of revealing the actual correlations among cognitive abilities and brain structure in the population. However, a drawback to this approach is that doing so could also amplify these associations, confounding our findings. On the other hand, regressing out age might enable us to detect correlations *beyond* age, possibility revealing core relations among variables independent of stereotypical neurocognitive development (e.g., older participants normally score higher on cognitive tasks and have larger brains as they mature). However, this might also remove developmental associations of interest (e.g., age may function as a moderator of cognitive and neural growth).

Here we compare the partial correlations matrices for the two analysis paths (age node used in network estimation vs. age node regressed out before estimation) for both single and multilayer networks (Supplementary Table 2). This analysis demonstrates that, regardless of how age is accounted for in estimation, the partial correlation networks are very similar to each other.

**Supplementary Table 2.** Comparisons between partial correlation networks (age included in estimation vs age regressed out before estimation). These include summary statistics such as mean, (standard deviation), [range] and Pearson correlations between PC graph weights using pairwise complete observations to account for missingness.

| **Network Type** | **Age Included**<br>**in Estimation** | **Age Regressed Out**<br>**before Estimation** | **Pearson Correlation** |
| --- | --- | --- | --- |
| Cognitive | 0.08 (0.11)<br>[0, 0.63] | 0.08 (0.12)<br>[0, 0.65] | 0.98 |
| Grey Matter | 0.09 (0.14)<br>[-0.15, 0.52] | 0.09 (0.14)<br>[-0.15, 0.52] | 1(rounded from 0.999) |
| White Matter | 0.08 (0.11)<br>[0, 0.44] | 0.08 (0.13)<br>[-0.2, 0.49] | 0.93 |
| Cognitive-grey matter | 0.04 (0.1)<br>[-0.12, 0.64] | 0.03 (0.1)<br>[-0.14, 0.65] | 0.94 |
| Cognitive-white matter | 0.04 (0.1)<br>[-0.2, 0.65] | 0.03 (0.1)<br>[-0.24, 0.66] | 0.94 |
| Tri-layer | 0.02 (0.08)<br>[-0.2, 0.66] | 0.02 (0.07)<br>[0, 0.64] | 0.88 |

### Teasing Apart the Relations of Cortical Volume to General Intelligence: Multilayer Analysis Using Cortical Surface Area and Thickness

Lastly, we partitioned cortical volume into its constituent parts, cortical surface area and thickness, to compare their partial correlations and community structures when combined with white matter and general intelligence (Supplementary Figures 4 and 5). Although not conclusive, the effect seen for cortical volume in the main manuscript appears to be driven by cortical surface area, but not thickness, as exhibited by greater inter-connectivity among domains (brain vs behavior). Finally, bridge strength showed the same pattern as in the main manuscript, except for the cortical thickness tri-layer network, where white matter appears to dominate the bridge strength centrality (Supplementary Figure 5).

**Supplementary Figure 4.**
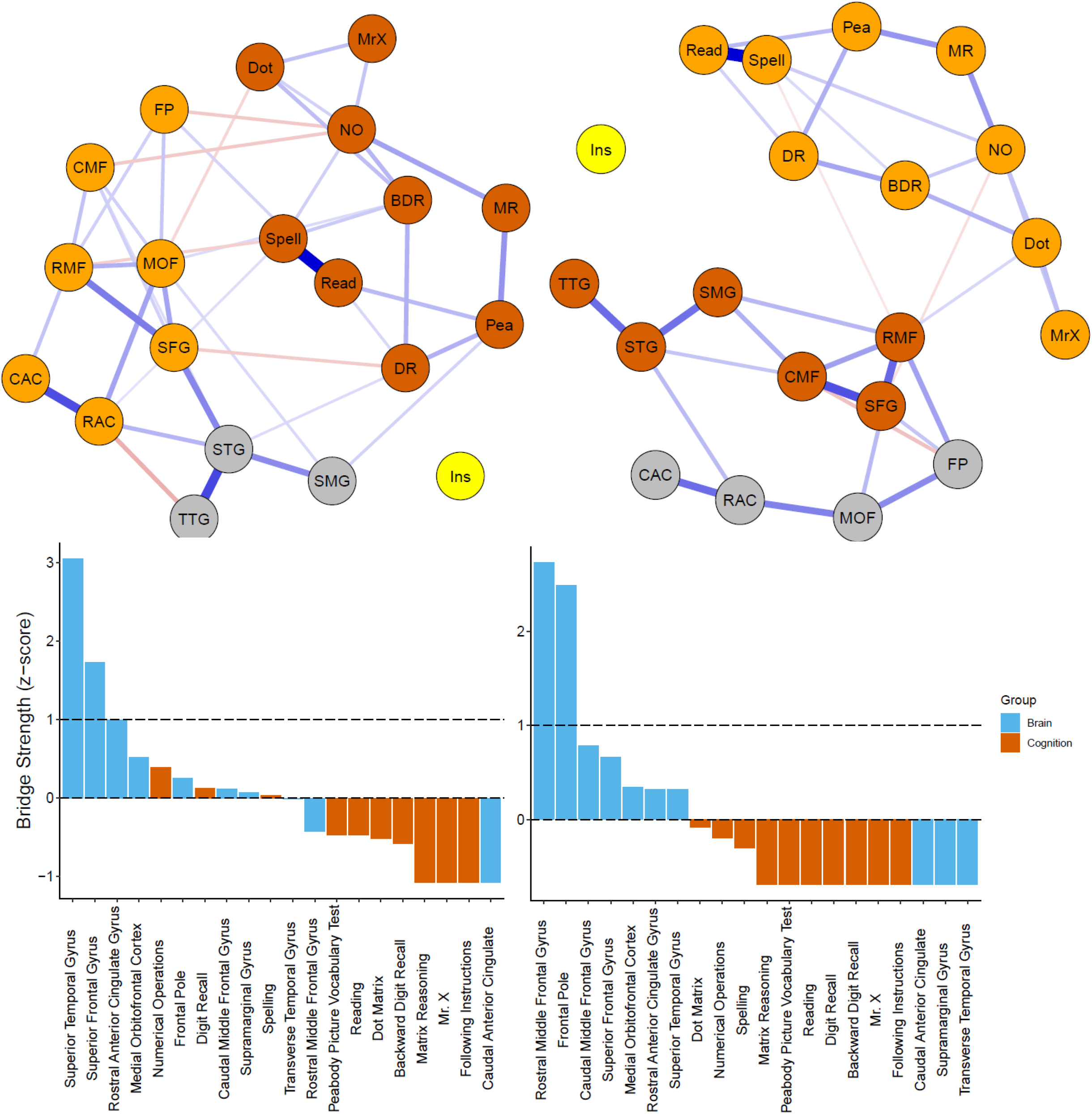
Top: Network visualizations (spring layout) of partial correlation CALM bi-layer grey matter (surface area (left) and cortical thickness (right)) networks. Nodes are grouped according to Walktrap algorithm results (see above). Bottom: Bridge centrality estimates (z-scores) for CALM bi-layer grey matter (surface area (left) and cortical thickness (right)) networks. Dashed lines indicate mean strength and one standard deviation above the mean.

**Supplementary Figure 5.**
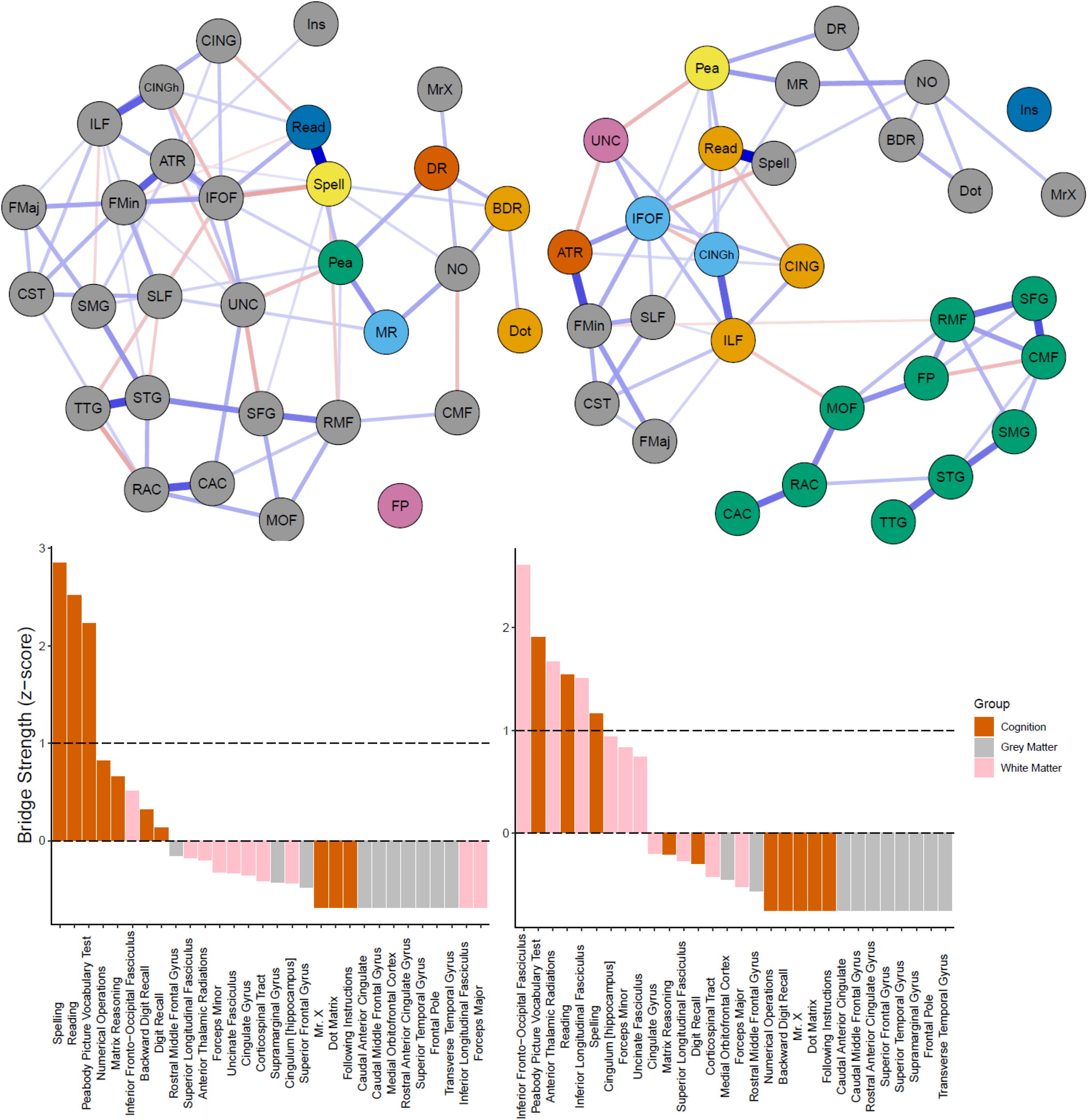
Top: Network visualizations (spring layout) of partial correlation CALM tri-layer grey matter (surface area (left) and cortical thickness (right)) networks. Nodes are grouped according to Walktrap algorithm results (see above). Bottom: Bridge centrality estimates (z-scores) for CALM tri-layer grey matter (surface area (left) and cortical thickness (right)) networks. Dashed lines indicate mean strength and one standard deviation above the mean.

## Footnotes

1 Dworkin, Jordan D., et al. “The extent and drivers of gender imbalance in neuroscience reference lists.” *Nature neuroscience* 23.8 (2020): 918-926.

2 Fulvio, Jacqueline M., Ileri Akinnola, and Bradley R. Postle. “Gender (im) balance in citation practices in cognitive neuroscience.” *Journal of Cognitive Neuroscience* 33.1 (2021): 3-7.

3 Bertolero, Maxwell A., et al. “Racial and ethnic imbalance in neuroscience reference lists and intersections with gender.” *BioRxiv* (2020).

## REFERENCES

Akarca, Danyal, Petra E. Vértes, Edward T. Bullmore, the CALM Team, and Duncan E. Astle. 2020. “A Generative Network Model of Neurodevelopment.” BioRxiv, August, 2020.08.13.249391. https://doi.org/10.1101/2020.08.13.249391.

Alexander-Bloch, Aaron, Jay N. Giedd, and Ed Bullmore. 2013. “Imaging Structural Co-Variance between Human Brain Regions.” Nature Reviews Neuroscience 14 (5): 322–36. https://doi.org/10.1038/nrn3465.

Alloway, Tracy P. 2007. “Automated Working Memory Assessment (AWMA).” London: Harcourt Assessment.

Amestoy et al., See AUTHORS file igraph author. 2020. Igraph: Network Analysis and Visualization (version 1.2.6). https://CRAN.R-project.org/package=igraph.

Avants, B. B., C. L. Epstein, M. Grossman, and J. C. Gee. 2008. “Symmetric Diffeomorphic Image Registration with Cross-Correlation: Evaluating Automated Labeling of Elderly and Neurodegenerative Brain.” Medical Image Analysis, Special Issue on The Third International Workshop on Biomedical Image Registration – WBIR 2006, 12 (1): 26–41. https://doi.org/10.1016/j.media.2007.06.004.

Barabási, Albert-László. 2016. Network Science. Cambridge University Press. http://networksciencebook.com/.

Barbey, Aron K. 2018. “Network Neuroscience Theory of Human Intelligence.” Trends in Cognitive Sciences 22 (1): 8–20. https://doi.org/10.1016/j.tics.2017.10.001.

Bassett, Danielle S., and Edward T. Bullmore. 2017. “Small-World Brain Networks Revisited.” The Neuroscientist 23 (5): 499–516. https://doi.org/10.1177/1073858416667720.

Bassett, Danielle S, and Olaf Sporns. 2017. “Network Neuroscience.” Nature Neuroscience 20 (3): 353–64. https://doi.org/10.1038/nn.4502.

Bassett, Danielle Smith, and Ed Bullmore. 2006. “Small-World Brain Networks.” The Neuroscientist 12 (6): 512–23. https://doi.org/10.1177/1073858406293182.

Basten, Ulrike, Kirsten Hilger, and Christian J. Fiebach. 2015. “Where Smart Brains Are Different: A Quantitative Meta-Analysis of Functional and Structural Brain Imaging Studies on Intelligence.” Intelligence 51 (July): 10–27. https://doi.org/10.1016/j.intell.2015.04.009.

Bathelt, Joe, Hilde M. Geurts, and Denny Borsboom. 2020. “More than the Sum of Its Parts: Merging Network Psychometrics and Network Neuroscience with Application in Autism.” BioRxiv, November, 2020.11.17.386276. https://doi.org/10.1101/2020.11.17.386276.

Bertolero, M. A., B. T. T. Yeo, and M. D’Esposito. 2017. “The Diverse Club.” Nature Communications 8 (1): 1277. https://doi.org/10.1038/s41467-017-01189-w.

Bertolero, Maxwell A., B. T. Thomas Yeo, Danielle S. Bassett, and Mark D’Esposito. 2018. “A Mechanistic Model of Connector Hubs, Modularity and Cognition.” Nature Human Behaviour 2 (10): 765–77. https://doi.org/10.1038/s41562-018-0420-6.

Bianconi, Ginestra. 2018. Multilayer Networks: Structure and Function. Oxford University Press.

Borsboom, Denny. 2017. “A Network Theory of Mental Disorders.” World Psychiatry 16 (1): 5–13. https://doi.org/10.1002/wps.20375.

Bullmore, Ed, and Olaf Sporns. 2012. “The Economy of Brain Network Organization.” Nature Reviews Neuroscience 13 (5): 336–49. https://doi.org/10.1038/nrn3214.

Calvin, C. M., I. J. Deary, C. Fenton, B. A. Roberts, G. Der, N. Leckenby, and G. D. Batty. 2011. “Intelligence in Youth and All-Cause-Mortality: Systematic Review with Meta-Analysis.” International Journal of Epidemiology 40 (3): 626–44. https://doi.org/10.1093/ije/dyq190.

Carroll, John B. 1993. Human Cognitive Abilities: A Survey of Factor-Analytic Studies. Cambridge University Press.

Casey, B. J., Tariq Cannonier, May I. Conley, Alexandra O. Cohen, Deanna M. Barch, Mary M. Heitzeg, Mary E. Soules, et al. 2018. “The Adolescent Brain Cognitive Development (ABCD) Study: Imaging Acquisition across 21 Sites.” Developmental Cognitive Neuroscience, The Adolescent Brain Cognitive Development (ABCD) Consortium: Rationale, Aims, and Assessment Strategy, 32 (August): 43–54. https://doi.org/10.1016/j.dcn.2018.03.001.

Cattell, Raymond B. 1971. Abilities: Their Structure, Growth, and Action. Abilities: Their Structure, Growth, and Action. Oxford, England: Houghton Mifflin.

Dablander, Fabian, and Max Hinne. 2019. “Node Centrality Measures Are a Poor Substitute for Causal Inference.” Scientific Reports 9 (1): 1–13. https://doi.org/10.1038/s41598-019-43033-9.

Dale, Anders M., Bruce Fischl, and Martin I. Sereno. 1999. “Cortical Surface-Based Analysis: I. Segmentation and Surface Reconstruction.” NeuroImage 9 (2): 179–94. https://doi.org/10.1006/nimg.1998.0395.

Deary, Ian J., Martha C. Whiteman, John M. Starr, Lawrence J. Whalley, and Helen C. Fox. 2004. “The Impact of Childhood Intelligence on Later Life: Following Up the Scottish Mental Surveys of 1932 and 1947.” Journal of Personality and Social Psychology 86 (1): 130–47. https://doi.org/10.1037/0022-3514.86.1.130.

Desikan, Rahul S., Florent Ségonne, Bruce Fischl, Brian T. Quinn, Bradford C. Dickerson, Deborah Blacker, Randy L. Buckner, et al. 2006. “An Automated Labeling System for Subdividing the Human Cerebral Cortex on MRI Scans into Gyral Based Regions of Interest.” NeuroImage 31 (3): 968–80. https://doi.org/10.1016/j.neuroimage.2006.01.021.

Dunn, Lloyd M., and Douglas M. Dunn. 2007. “PPVT-4: Peabody Picture Vocabulary Test.” Pearson Assessments.

Epskamp, Sacha, Denny Borsboom, and Eiko I. Fried. 2018. “Estimating Psychological Networks and Their Accuracy: A Tutorial Paper.” Behavior Research Methods 50 (1): 195–212. https://doi.org/10.3758/s13428-017-0862-1.

Epskamp, Sacha, Giulio Costantini, Jonas Haslbeck, Adela Isvoranu, Angelique O. J. Cramer, Lourens J. Waldorp, Verena D. Schmittmann, and Denny Borsboom. 2020. Qgraph: Graph Plotting Methods, Psychometric Data Visualization and Graphical Model Estimation (version 1.6.5). https://CRAN.R-project.org/package=qgraph.

Epskamp, Sacha, and Eiko I. Fried. 2018. “A Tutorial on Regularized Partial Correlation Networks.” Psychological Methods 23 (4): 617–34. https://doi.org/10.1037/met0000167.

Epskamp, Sacha, and Eiko I. Fried. 2020. Bootnet: Bootstrap Methods for Various Network Estimation Routines (version 1.4.3). https://CRAN.R-project.org/package=bootnet.

Epskamp, Sacha, Gunter Maris, Lourens J. Waldorp, and Denny Borsboom. 2018. “Network Psychometrics.” In The Wiley Handbook of Psychometric Testing, 953–86. John Wiley & Sons, Ltd. https://doi.org/10.1002/9781118489772.ch30.

Fischl, B., M. I. Sereno, and A. M. Dale. 1999. “Cortical Surface-Based Analysis. II: Inflation, Flattening, and a Surface-Based Coordinate System.” NeuroImage 9 (2): 195–207. https://doi.org/10.1006/nimg.1998.0396.

Fischl, Bruce, and Anders M. Dale. 2000. “Measuring the Thickness of the Human Cerebral Cortex from Magnetic Resonance Images.” Proceedings of the National Academy of Sciences 97 (20): 11050–55. https://doi.org/10.1073/pnas.200033797.

Fischl, Bruce, David H. Salat, Evelina Busa, Marilyn Albert, Megan Dieterich, Christian Haselgrove, Andre van der Kouwe, et al. 2002. “Whole Brain Segmentation: Automated Labeling of Neuroanatomical Structures in the Human Brain.” Neuron 33 (3): 341–55. https://doi.org/10.1016/S0896-6273(02)00569-X.

Fornito, Alex, and Edward T. Bullmore. 2012. “Connectomic Intermediate Phenotypes for Psychiatric Disorders.” Frontiers in Psychiatry 3. https://doi.org/10.3389/fpsyt.2012.00032.

Fornito, Alex, Andrew Zalesky, and Edward Bullmore. 2016. Fundamentals of Brain Network Analysis. Academic Press.

Fried, Eiko I. 2020a. “Lack of Theory Building and Testing Impedes Progress in the Factor and Network Literature,” January. https://doi.org/10.17605/OSF.IO/6CTS9.

Fried, Eiko I. 2020b. “Lack of Theory Building and Testing Impedes Progress in The Factor and Network Literature.” Psychological Inquiry 31 (4): 271–88. https://doi.org/10.1080/1047840X.2020.1853461.

Fried, Eiko I., and Angélique O. J. Cramer. 2017. “Moving Forward: Challenges and Directions for Psychopathological Network Theory and Methodology.” Perspectives on Psychological Science 12 (6): 999–1020. https://doi.org/10.1177/1745691617705892.

Fried, Eiko I., and Donald J. Robinaugh. 2020. “Systems All the Way down: Embracing Complexity in Mental Health Research.” BMC Medicine 18 (1): 205. https://doi.org/10.1186/s12916-020-01668-w.

Friedman, Jerome, Trevor Hastie, and Robert Tibshirani. 2008. “Sparse Inverse Covariance Estimation with the Graphical Lasso.” Biostatistics 9 (3): 432–41. https://doi.org/10.1093/biostatistics/kxm045.

Gates, Kathleen M., Teague Henry, Doug Steinley, and Damien A. Fair. 2016. “A Monte Carlo Evaluation of Weighted Community Detection Algorithms.” Frontiers in Neuroinformatics 10. https://doi.org/10.3389/fninf.2016.00045.

Gathercole, Susan E., Emily Durling, Matthew Evans, Sarah Jeffcock, and Sarah Stone. 2008. “Working Memory Abilities and Children’s Performance in Laboratory Analogues of Classroom Activities.” Applied Cognitive Psychology 22 (8): 1019–37. https://doi.org/10.1002/acp.1407.

Gathercole, Susan E., Elisabet Service, Graham J. Hitch, Anne-Marie Adams, and Amanda J. Martin. 1999. “Phonological Short-Term Memory and Vocabulary Development: Further Evidence on the Nature of the Relationship.” Applied Cognitive Psychology 13 (1): 65–77. https://doi.org/10.1002/(SICI)1099-0720(199902)13:1<65::AID-ACP548>3.0.CO;2-O.

Girn, Manesh, Caitlin Mills, and Kalina Christoff. 2019. “Linking Brain Network Reconfiguration and Intelligence: Are We There Yet?” Trends in Neuroscience and Education 15 (June): 62–70. https://doi.org/10.1016/j.tine.2019.04.001.

Graham, Mark S., Ivana Drobnjak, and Hui Zhang. 2016. “Realistic Simulation of Artefacts in Diffusion MRI for Validating Post-Processing Correction Techniques.” NeuroImage 125 (January): 1079–94. https://doi.org/10.1016/j.neuroimage.2015.11.006.

Gu, Shi, Fabio Pasqualetti, Matthew Cieslak, Qawi K. Telesford, Alfred B. Yu, Ari E. Kahn, John D. Medaglia, et al. 2015. “Controllability of Structural Brain Networks.” Nature Communications 6 (1): 1–10. https://doi.org/10.1038/ncomms9414.

Hegelund, Emilie Rune, Trine Flensborg-Madsen, Jesper Dammeyer, and Erik Lykke Mortensen. 2018. “Low IQ as a Predictor of Unsuccessful Educational and Occupational Achievement: A Register-Based Study of 1,098,742 Men in Denmark 1968–2016.” Intelligence 71 (November): 46–53. https://doi.org/10.1016/j.intell.2018.10.002.

Henry, Teague R., Donald Robinaugh, and Eiko I. Fried. 2020. “On the Control of Psychological Networks.” PsyArXiv. https://doi.org/10.31234/osf.io/7vpz2.

Heuvel, Martijn P. van den, and Olaf Sporns. 2011. “Rich-Club Organization of the Human Connectome.” Journal of Neuroscience 31 (44): 15775–86. https://doi.org/10.1523/JNEUROSCI.3539-11.2011.

Hilland, Eva, Nils I. Landrø, Brage Kraft, Christian K. Tamnes, Eiko I. Fried, Luigi A. Maglanoc, and Rune Jonassen. 2020. “Exploring the Links between Specific Depression Symptoms and Brain Structure: A Network Study.” Psychiatry and Clinical Neurosciences 74 (3): 220–21. https://doi.org/10.1111/pcn.12969.

Holmes, Joni, Annie Bryant, Susan Elizabeth Gathercole, and the CALM Team. 2019. “Protocol for a Transdiagnostic Study of Children with Problems of Attention, Learning and Memory (CALM).” BMC Pediatrics 19 (10). https://doi.org/10.1186/s12887-018-1385-3.

Hua, Kegang, Jiangyang Zhang, Setsu Wakana, Hangyi Jiang, Xin Li, Daniel S. Reich, Peter A. Calabresi, James J. Pekar, Peter C.M. van Zijl, and Susumu Mori. 2008. “Tract Probability Maps in Stereotaxic Spaces: Analyses of White Matter Anatomy and Tract-Specific Quantification.” NeuroImage 39 (1): 336–47. https://doi.org/10.1016/j.neuroimage.2007.07.053.

Jones, Payton. 2020. Networktools: Tools for Identifying Important Nodes in Networks (version 1.2.3). https://CRAN.R-project.org/package=networktools.

Jones, Payton J., Ruofan Ma, and Richard J. McNally. 2019. “Bridge Centrality: A Network Approach to Understanding Comorbidity.” Multivariate Behavioral Research 0 (0): 1–15. https://doi.org/10.1080/00273171.2019.1614898.

Jung, Rex E., and Richard J. Haier. 2007. “The Parieto-Frontal Integration Theory (P-FIT) of Intelligence: Converging Neuroimaging Evidence.” Behavioral and Brain Sciences 30 (02): 135. https://doi.org/10.1017/S0140525X07001185.

Kan, Kees-Jan, Han L.J. van der Maas, and Stephen Z. Levine. 2019. “Extending Psychometric Network Analysis: Empirical Evidence against g in Favor of Mutualism?” Intelligence 73 (March): 52–62. https://doi.org/10.1016/j.intell.2018.12.004.

Khundrakpam, Budhachandra S., John D. Lewis, Andrew Reid, Sherif Karama, Lu Zhao, Francois Chouinard-Decorte, and Alan C. Evans. 2017. “Imaging Structural Covariance in the Development of Intelligence.” NeuroImage 144 (January): 227–40. https://doi.org/10.1016/j.neuroimage.2016.08.041.

Kievit, Rogier A., Simon W. Davis, John Griffiths, Marta M. Correia, Cam-CAN, and Richard N. Henson. 2016. “A Watershed Model of Individual Differences in Fluid Intelligence.” Neuropsychologia 91 (October): 186–98. https://doi.org/10.1016/j.neuropsychologia.2016.08.008.

Kievit, Rogier A., Delia Fuhrmann, Gesa Sophia Borgeest, Ivan L. Simpson-Kent, and Richard N. A. Henson. 2018. “The Neural Determinants of Age-Related Changes in Fluid Intelligence: A Pre-Registered, Longitudinal Analysis in UK Biobank.” Wellcome Open Research 3 (June). https://doi.org/10.12688/wellcomeopenres.14241.2.

Kievit, Rogier A., Abe D. Hofman, and Kate Nation. 2019. “Mutualistic Coupling Between Vocabulary and Reasoning in Young Children: A Replication and Extension of the Study by Kievit et al. (2017).” Psychological Science 30 (8): 1245–52. https://doi.org/10.1177/0956797619841265.

Kievit, Rogier A., Ulman Lindenberger, Ian M. Goodyer, Peter B. Jones, Peter Fonagy, Edward T. Bullmore, and Raymond J. Dolan. 2017. “Mutualistic Coupling Between Vocabulary and Reasoning Supports Cognitive Development During Late Adolescence and Early Adulthood.” Psychological Science 28 (10): 1419–31. https://doi.org/10.1177/0956797617710785.

Kievit, Rogier A, and Ivan L Simpson-Kent. 2021. “It’s About Time: Towards a Longitudinal Cognitive Neuroscience of Intelligence.” In The Cambridge Handbook of Intelligence and Cognitive Neuroscience. Cambridge University Press.

Kievit, Rogier, Willem Eduard Frankenhuis, Lourens Waldorp, and Denny Borsboom. 2013. “Simpson’s Paradox in Psychological Science: A Practical Guide.” Frontiers in Psychology 4. https://doi.org/10.3389/fpsyg.2013.00513.

Krogsrud, Stine K., Anders M. Fjell, Christian K. Tamnes, Håkon Grydeland, Paulina Due-Tønnessen, Atle Bjørnerud, Cassandra Sampaio-Baptista, Jesper Andersson, Heidi Johansen-Berg, and Kristine B. Walhovd. 2018. “Development of White Matter Microstructure in Relation to Verbal and Visuospatial Working Memory—A Longitudinal Study.” Edited by Hao Huang. PLOS ONE 13 (4): e0195540. https://doi.org/10.1371/journal.pone.0195540.

Levenstein, Daniel, Veronica A. Alvarez, Asohan Amarasingham, Habiba Azab, Richard C. Gerkin, Andrea Hasenstaub, Ramakrishnan Iyer, et al. 2020. “On the Role of Theory and Modeling in Neuroscience.” ArXiv:2003.13825 [q-Bio], April. http://arxiv.org/abs/2003.13825.

Levine, Stephen Z., and Stefan Leucht. 2016. “Identifying a System of Predominant Negative Symptoms: Network Analysis of Three Randomized Clinical Trials.” Schizophrenia Research 178 (1): 17–22. https://doi.org/10.1016/j.schres.2016.09.002.

Maas, Han L. J. van der, Conor V. Dolan, Raoul P. P. P. Grasman, Jelte M. Wicherts, Hilde M. Huizenga, and Maartje E. J. Raijmakers. 2006. “A Dynamical Model of General Intelligence: The Positive Manifold of Intelligence by Mutualism.” Psychological Review 113 (4): 842–61. https://doi.org/10.1037/0033-295X.113.4.842.

Maas, Han L. J. van der, Kees-Jan Kan, Maarten Marsman, and Claire E. Stevenson. 2017. “Network Models for Cognitive Development and Intelligence.” Journal of Intelligence 5 (2): 16. https://doi.org/10.3390/jintelligence5020016.

Mareva, Silvana, and Joni Holmes. 2020. “Network Models of Learning and Cognition in Typical and Atypical Learners,” September. https://doi.org/10.31219/osf.io/3tn5m.

Marr, D., and T. Poggio. 1976. “From Understanding Computation to Understanding Neural Circuitry,” May. https://dspace.mit.edu/handle/1721.1/5782.

Meunier, David, Renaud Lambiotte, and Edward T. Bullmore. 2010. “Modular and Hierarchically Modular Organization of Brain Networks.” Frontiers in Neuroscience 4. https://doi.org/10.3389/fnins.2010.00200.

Mooij, Susanne M.M. de, Richard N.A. Henson, Lourens J. Waldorp, and Rogier A. Kievit. 2018. “Age Differentiation within Gray Matter, White Matter, and between Memory and White Matter in an Adult Life Span Cohort.” The Journal of Neuroscience 38 (25): 5826–36. https://doi.org/10.1523/JNEUROSCI.1627-17.2018.

Navas-Sánchez, Francisco J., Yasser Alemán-Gómez, Javier Sánchez-Gonzalez, Juan A. Guzmán-De-Villoria, Carolina Franco, Olalla Robles, Celso Arango, and Manuel Desco. 2014. “White Matter Microstructure Correlates of Mathematical Giftedness and Intelligence Quotient: White Matter Microstructure.” Human Brain Mapping 35 (6): 2619–31. https://doi.org/10.1002/hbm.22355.

Newman, M. E. J. 2006. “Modularity and Community Structure in Networks.” Proceedings of the National Academy of Sciences 103 (23): 8577–82. https://doi.org/10.1073/pnas.0601602103.

Peng, Peng, Marcia Barnes, CuiCui Wang, Wei Wang, Shan Li, H. Lee Swanson, William Dardick, and Sha Tao. 2018. “A Meta-Analysis on the Relation between Reading and Working Memory.” Psychological Bulletin 144 (1): 48–76. https://doi.org/10.1037/bul0000124.

Pons, Pascal, and Matthieu Latapy. 2005. “Computing Communities in Large Networks Using Random Walks.” In Computer and Information Sciences - ISCIS 2005, edited by pInar Yolum, Tunga Güngör, Fikret Gürgen, and Can Özturan, 284–93. *Lecture Notes in Computer Science*. Berlin, Heidelberg: Springer. https://doi.org/10.1007/11569596_31.

R Core Team. 2020. “R: A Language and Environment for Statistical Computing. R Foundation for Statistical Computing, Vienna.” https://www.R-project.org.

Reid, Andrew T., Drew B. Headley, Ravi D. Mill, Ruben Sanchez-Romero, Lucina Q. Uddin, Daniele Marinazzo, Daniel J. Lurie, et al. 2019. “Advancing Functional Connectivity Research from Association to Causation.” Nature Neuroscience 22 (11): 1751–60. https://doi.org/10.1038/s41593-019-0510-4.

Rhemtulla, Mijke, Riet van Bork, and A. O. J. Cramer. 2020. “Cross-Lagged Network Models.” Multivariate Behavioral Research. https://research.tilburguniversity.edu/en/publications/cross-lagged-network-models.

Robinaugh, Donald J., Ria H. A. Hoekstra, Emma R. Toner, and Denny Borsboom. 2019. “The Network Approach to Psychopathology: A Review of the Literature 2008–2018 and an Agenda for Future Research.” Psychological Medicine, 1–14. https://doi.org/10.1017/S0033291719003404.

Rohrer, Julia M. 2018. “Thinking Clearly About Correlations and Causation: Graphical Causal Models for Observational Data.” Advances in Methods and Practices in Psychological Science 1 (1): 27–42. https://doi.org/10.1177/2515245917745629.

Schmank, Christopher J., Sara Anne Goring, Kristof Kovacs, and Andrew R. A. Conway. 2019. “Psychometric Network Analysis of the Hungarian WAIS.” Journal of Intelligence 7 (3): 21. https://doi.org/10.3390/jintelligence7030021.

Schmiedek, Florian, Martin Lövdén, Timo von Oertzen, and Ulman Lindenberger. 2020. “Within-Person Structures of Daily Cognitive Performance Differ from between-Person Structures of Cognitive Abilities.” PeerJ 8 (June): e9290. https://doi.org/10.7717/peerj.9290.

Seidlitz, Jakob, František Váša, Maxwell Shinn, Rafael Romero-Garcia, Kirstie J. Whitaker, Petra E. Vértes, Konrad Wagstyl, et al. 2018. “Morphometric Similarity Networks Detect Microscale Cortical Organization and Predict Inter-Individual Cognitive Variation.” Neuron 97 (1): 231–247.e7. https://doi.org/10.1016/j.neuron.2017.11.039.

Simpson-Kent, Ivan L., Delia Fuhrmann, Joe Bathelt, Jascha Achterberg, Gesa Sophia Borgeest, and Rogier A. Kievit. 2020. “Neurocognitive Reorganization between Crystallized Intelligence, Fluid Intelligence and White Matter Microstructure in Two Age-Heterogeneous Developmental Cohorts.” Developmental Cognitive Neuroscience 41 (February): 100743. https://doi.org/10.1016/j.dcn.2019.100743.

Smith, Stephen M. 2002. “Fast Robust Automated Brain Extraction.” Human Brain Mapping 17 (3): 143–55. https://doi.org/10.1002/hbm.10062.

Solé-Casals, Jordi, Josep M. Serra-Grabulosa, Rafael Romero-Garcia, Gemma Vilaseca, Ana Adan, Núria Vilaró, Núria Bargalló, and Edward T. Bullmore. 2019. “Structural Brain Network of Gifted Children Has a More Integrated and Versatile Topology.” Brain Structure and Function 224 (7): 2373–83. https://doi.org/10.1007/s00429-019-01914-9.

Spearman, Charles. 1904. “‘General Intelligence,’ Objectively Determined and Measured.” The American Journal of Psychology 15 (2): 201–92.

Sporns, Olaf, and Richard F. Betzel. 2016. “Modular Brain Networks.” Annual Review of Psychology 67 (January): 613–40. https://doi.org/10.1146/annurev-psych-122414-033634.

Wandell, Brian A. 2016. “Clarifying Human White Matter.” Annual Review of Neuroscience 39 (1): 103–28. https://doi.org/10.1146/annurev-neuro-070815-013815.

Wechsler, David. 2005. “Wechsler Individual Achievement Test-Second UK Edition (WIAT-II).” Pearson: London, UK.

Wechsler, David. 2008. “Wechsler Adult Intelligence Scale–Fourth Edition (WAIS–IV).” The Psychological Corporation, San Antonio, TX.

Wechsler, David. 2011. “Wechsler Abbreviated Scales of Intelligence-Second Edition (WASI-II).” Pearson: London, UK.

Zhang, Shuai, and R. Malatesha Joshi. 2020. “Longitudinal Relations between Verbal Working Memory and Reading in Students from Diverse Linguistic Backgrounds.” Journal of Experimental Child Psychology 190 (February): 104727. https://doi.org/10.1016/j.jecp.2019.104727.

